# Detection of ESKAPE pathogens and *Clostridioides difficile* in Simulated Skin Transmission Events with Metagenomic and Metatranscriptomic Sequencing

**DOI:** 10.1101/2021.03.04.433847

**Authors:** Krista L. Ternus, Nicolette C. Keplinger, Anthony D. Kappell, Gene D. Godbold, Veena Palsikar, Carlos A. Acevedo, Katharina L. Weber, Danielle S. LeSassier, Kathleen Q. Schulte, Nicole M. Westfall, F. Curtis Hewitt

**Affiliations:** Signature Science, LLC, 8329 North Mopac Expressway, Austin, Texas, USA; Signature Science, LLC, 1670 Discovery Drive, Charlottesville, VA, USA

**Keywords:** Metagenomics, Metatranscriptomics, ESKAPE Pathogens, *Clostridioides difficile*, Antibiotic Resistance, Epidemiology, Bioinformatics

## Abstract

1

**Background:** Antimicrobial resistance is a significant global threat, posing major public health risks and economic costs to healthcare systems. Bacterial cultures are typically used to diagnose healthcare-acquired infections (HAI); however, culture-dependent methods provide limited presence/absence information and are not applicable to all pathogens. Next generation sequencing (NGS) has the capacity to detect a wide variety of pathogens, virulence elements, and antimicrobial resistance (AMR) signatures in healthcare settings without the need for culturing, but few research studies have explored how NGS could be used to detect viable human pathogen transmission events under different HAI-relevant scenarios.

**Methods:** The objective of this project was to assess the capability of NGS-based methods to detect the direct and indirect transmission of high priority healthcare-related pathogens. DNA was extracted and sequenced from a previously published study exploring pathogen transfer with simulated skin containing background microorganisms, which allowed for complementary culture and metagenomic analysis comparisons. RNA was also isolated from an additional set of samples to evaluate metatranscriptomic analysis methods at different concentrations.

**Results:** Using various analysis methods and custom reference databases, both pathogenic and non-pathogenic members of the microbial community were taxonomically identified. Virulence and AMR genes known to reside within the community were also routinely detected. Ultimately, pathogen abundance within the overall microbial community played the largest role in successful taxonomic classification and gene identification.

**Conclusions:** These results illustrate the utility of metagenomic analysis in clinical settings or for epidemiological studies, but also highlight the limits associated with the detection and characterization of pathogens at low abundance in a microbial community.

## 3 Introduction

The estimated number of annual deaths due to infections from multidrug resistant organisms is upwards of ∼70,000 for individuals within inpatient hospital care and ∼80,000 for those in outpatient care in the United States, based on 2010 mortality rates (Burnham et al., 2019). The ESKAPE pathogens, consisting of *Enterococcus faecium, Staphylococcus aureus, Klebsiella pneumoniae, Acinetobacter baumannii, Pseudomonas aeruginosa*, and *Enterobacter species* (Boucher et al., 2009), are responsible for many drug-resistant healthcare-acquired infections (HAIs) (Boucher et al., 2009; Santajit and Indrawattana, 2016). Along with *Clostridioides difficile* (Slimings and Riley, 2014), these pathogens are the leading causes of nosocomial infections (Boucher et al., 2009; Santajit and Indrawattana, 2016). Culture-based methods within clinical laboratories are typically utilized to identify and track HAI transmission, such as the nosocomial infections caused by ESKAPE pathogens and *C. difficile* (ESKAPE+C), (Didelot et al., 2012), but cultures have multiple drawbacks. Dead or unculturable pathogens will be overlooked by culture-dependent methods, even though usable biochemical signatures (e.g., DNA) persist. Culturing is primarily a method for identifying viable pathogens amenable to growth under certain conditions, aiming to confirm the presence of known pathogens at the species level. Once a putative pathogen species has been identified, multiple rounds of culturing and biochemical assays may be necessary to further characterize pathogens at the strain level or to identify antibiotic resistance activity.

Metagenomic and metatranscriptomic analyses of samples collected in a healthcare setting provide compelling alternatives to traditional culture-based pathogen identification. These analyses do not require pathogen viability or culturability; instead, collected cells are lysed and the nucleic acids are collected for sequencing. These approaches permit species or even strain level identifications of pathogens present within a sample without multiple rounds of culture analysis. Perhaps most importantly, sequencing approaches can provide valuable insights into gene content and expression, identifying components of the resistome and elements contributing to virulence in a clinical sample. Previous studies have evaluated the relationship between culture and metagenomic analysis, highlighting both successes and challenges for this technology (Didelot et al., 2012). Challenges of unbiased metagenomic or metatranscriptomic sequencing methods include complexities in developing standardized analysis protocols and databases, and pathogen concentrations falling below the limit of detection in relation to other organisms in the sample. In the current study, we constructed customized databases based on the known mock microbial community genome and gene content to explore the impact of different ESKAPE+C concentration levels and simulated HAI transfer scenarios on pathogen detection from metagenomic and metatranscriptomic sequence data.

Our research expands upon previously published data from a study establishing an *in vitro* method to model ESKAPE+C transmission using a synthetic skin surrogate (Weber et al., 2020). This prior study enabled the investigation of both direct (skin-to-skin) and indirect (skin-to fomite-to skin) pathogen transmission scenarios using VITRO SKIN^®^ N-19 to mimic human skin, including a simulated commensal skin flora (Figure 1). The commensal skin flora was included on both the pre-transfer and post-transfer coupon to simulate pathogen transfer from skin containing a mix of pathogen and commensal organisms to a second piece of skin containing only the existing commensal community. Different transfer scenarios of ESKAPE+C species, including multiple wash or decontamination steps and high or low spike-in concentrations, were evaluated using culture analysis. Additionally, nucleic acids were extracted from all sample replicates to compare sequence data with culture results. The resulting sample set had a wide range of relative pathogen abundance in comparison to the commensal community, which was ideal for evaluating metagenomic and metatranscriptomic analysis methods. Here, we present the results, contrasting the utility of metagenomic and metatranscriptomic analysis across a range of pathogen abundance within simulated clinical samples.

**Figure 1.**
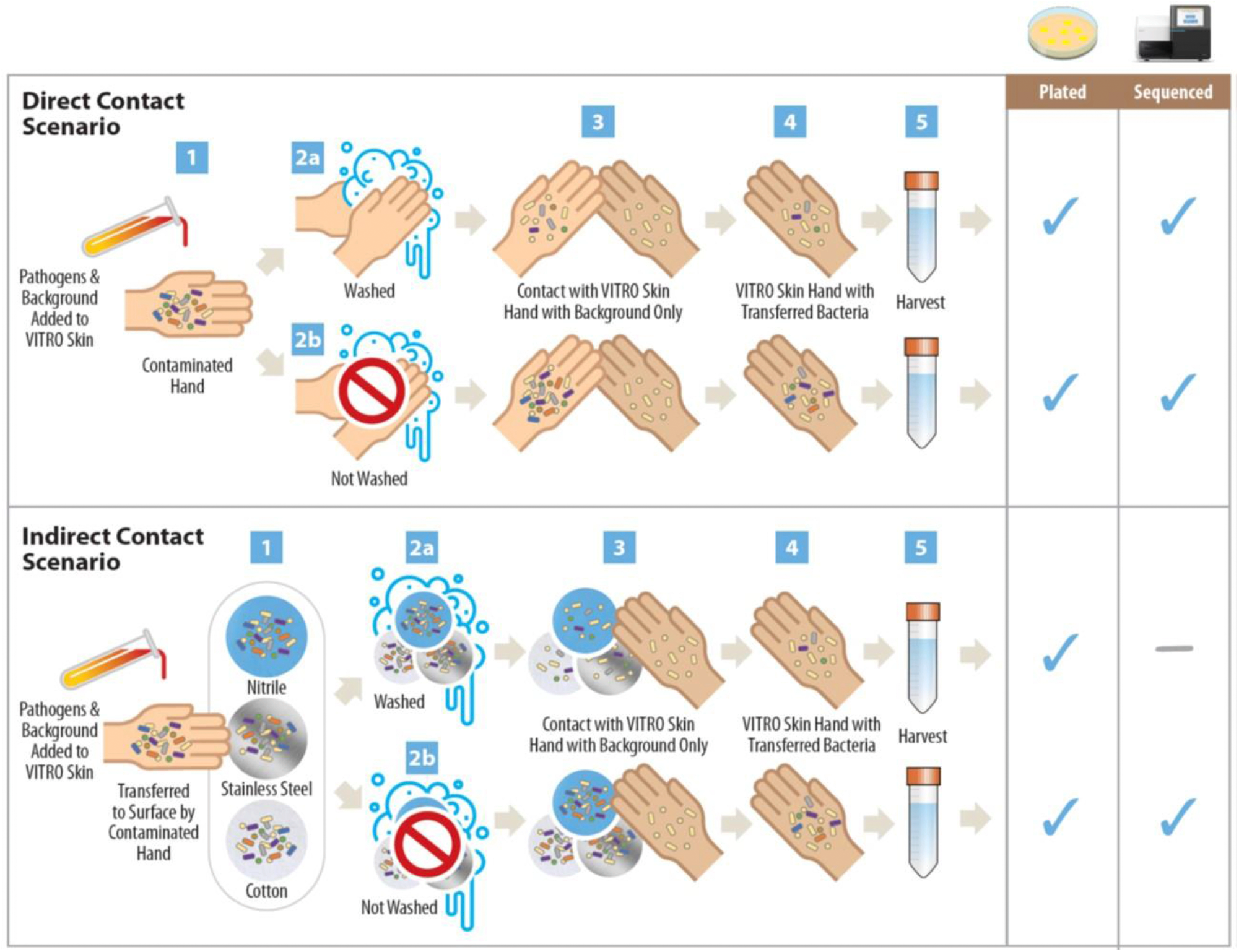
Sequences Collected from Direct and Indirect Contact Scenarios. For the direct contact scenario, the primary VITRO-SKIN^®^ coupon was inoculated with a mix of pathogen and background bacteria, representing a contaminated patient hand (step 1). The inoculated VITRO-SKIN^®^ was either washed (step 2a) or not washed (step 2b) and then briefly touched to a new, secondary VITRO-SKIN^®^ with only background microorganisms (step 3), simulating the touch transfer of bacteria from a sick patient to a clean healthcare worker. The secondary VITRO-SKIN^®^ representing the contaminated health care worker was then harvested (step 4) for culture and sequence analysis (step 5). Alternately, for the indirect contact scenario, the primary VITRO-SKIN^®^ coupon was inoculated with a mix of pathogen and background bacteria and touched to a surface of nitrile, stainless steel, or cotton, simulating bacterial transfer to the fomite in a healthcare setting (step 2c). A secondary VITRO-SKIN^®^ representing a clean health worker hand was then touched to the fomite and harvested as described above. Although our previous study analyzed culture data from the washed indirect transfer events (step 2b), those samples were not sequenced because the amount of available nucleic acid was anticipated to be too low for pathogen detection (Weber et al., 2020).

## 4 Materials and Methods

### 4.1 Bacterial Isolates and Sequence Data

Microorganisms used for this effort were sourced from American Type Culture Collection (ATCC) or the Centers for Disease Control and Prevention (CDC) Antimicrobial Resistance (AR) Isolate Bank as described previously (Weber et al., 2020). Any isolates that did not have existing published whole genome sequencing reference data were sequenced internally on an Illumina MiSeq^®^ FGx System. The new isolate sequence data produced by this study included *Enterococcus faecium* (CDC AR Bank #0579), *Clostridioides difficile* (ATCC 43598), *Brevibacterium linens* (ATCC 9172), *Corynebacterium matruchotii* (ATCC 14265), *Cutibacterium acnes* (ATCC 11827), *Escherichia coli* (ATCC 9637), *Lactobacillus gasseri* (ATCC 33323), *Micrococcus luteus* (ATCC 4698), *Staphylococcus epidermidis* (ATCC 12228), and *Streptococcus pyogenes* (ATCC 19615). All raw FASTQ sequences were submitted to NCBI SRA and subsequently processed for further analysis in this study. This included pre-processing to ensure a high quality of reads, mapping to reference genomes with 21x – 352x average genome coverage, and undergoing downstream assembly, gene identification, and taxonomic classification (Supplementary Tables 1-6).

Methods for bacterial culture, laboratory mixes, VITRO-SKIN^®^ coupon preparation for transfer experiments, and analysis of the culture data were previously published (Weber et al., 2020). Briefly, VITRO-SKIN^®^ N-19 (IMS Inc.) coupons were inoculated with “background organisms” to simulate a commensal skin microbial community at a constant concentration, and ESKAPE+C pathogens were added at two different high and low concentrations. Coupons were then allowed to dry before being touched to a second coupon containing only the commensal organisms, followed by a simulated handwashing step or non-handwashing step. Coupons were also transferred to a fomite surface (i.e., cotton, nitrile, stainless steel coupon), which was then subject to washing, decontamination, or no treatment before transfer to a skin coupon containing only commensal organisms.

### 4.2 Nucleic Acid Extractions

Each sample from the previous study was split, with one fraction used for culture analysis and the remaining sample used for nucleic acid extractions (Weber et al., 2020). Bacteria suspended in phosphate buffered saline (PBS) recovery buffer were transferred into 15 mL conical tubes and centrifuged at 5,000 x *g* for 20 minutes. The supernatant was removed, and DNA was extracted from the cell pellet using the ZymoBIOMICS DNA/RNA Miniprep Kit per manufacturer instructions. Total DNA yield was quantified using the Qubit™ dsDNA BR Assay Kit (Thermo Fisher Scientific) or Qubit™ dsDNA HS Assay Kit (Thermo Fisher Scientific) as appropriate per manufacturer instructions.

For the RNA control mixtures, a culture mixture was prepared that contained equal amounts (∼10^6^ CFU/mL) of all background bacteria and pathogens, representing the high pathogen concentrations for subsequent transfer events. Additional control of culture mixtures of all pathogen and background bacteria were made by titrating in equal amounts of each pathogen at ∼10^2^, ∼10^4^, or ∼10^6^ CFU/mL with a consistent amount of background bacteria at equal amounts (∼10^6^ CFU/mL) to mimic relative abundances commonly reported from the human hand in healthcare settings (WHO Guidelines on Hand Hygiene in Health Care: First Global Patient Safety Challenge Clean Care Is Safer Care, 2009). Samples were centrifuged at 5,000 x *g* for 20 minutes. The supernatant was removed, and RNA was extracted from the cell pellet using the ZymoBIOMICS DNA/RNA Miniprep Kit as per manufacturer instructions. Total RNA yield was quantified using the Qubit™ RNA HS Assay Kit (Thermo Fisher Scientific) as per manufacturer instructions.

### 4.3 RNA Preparation and cDNA Conversion

All RNA samples were converted to cDNA for subsequent library preparation and sequencing. First, sample mRNA was enriched using the MICROBExpress™ Bacterial mRNA Enrichment Kit (Thermo Fisher Scientific), followed by additional sample clean-up using the MEGAclear™ Transcription Clean-Up Kit (Thermo Fisher Scientific). Samples were then converted to cDNA using the SuperScript™ Double-Stranded cDNA Synthesis Kit (Thermo Fisher Scientific). All kits were used according to the manufacturer instructions. After each process, nucleic acid quantification was measured using either Qubit™ RNA HS Assay Kit or Qubit™ dsDNA HS Assay Kit (Thermo Fisher Scientific), as appropriate.

### 4.4 Library Preparation and Sequencing

Prior to library preparation, all samples (DNA for genomic analysis or cDNA for transcriptomic analysis) were diluted to 0.2 ng/µL based on measured Qubit™ values. For samples below 0.2 ng/µL, no dilution was performed. All samples were prepared for sequencing using the Nextera^®^ XT DNA Library Prep Kit (Illumina) as per the manufacturer’s protocol using AMPure XP (Beckman Coulter) bead-based normalization. Samples were diluted and denatured based on the recommended Nextera^®^ XT DNA bead-based normalization loading concentration. PhiX control (Illumina) was added at a final concentration of 1% to the denatured library, and libraries were loaded on a MiSeq^®^ Reagent v3 (Illumina) cartridge for sequencing. Sequencing was performed on a MiSeq^®^ FGx System (Illumina) in research use only (RUO) mode using the MiSeq^®^ Reagent Kit v3 (Illumina) with paired-end read lengths of 75 base pairs and results generated in FASTQ files. A bioinformatics quality control pipeline verified that all FASTQ files were of sufficient quality for downstream processing and analysis.

### 4.5 Bioinformatics Analysis

All bioinformatics tools and databases used in this analysis were open source, and more information, including the commands used with each of the bioinformatics tools is available in Supplementary Table 7. Quality control for sequence data analysis was performed with FastQC (Andrews, S, 2010), Trimmomatic (Bolger et al., 2014), and MultiQC (Ewels et al., 2016). Mash distance was used to identify the closest available reference genome with the trimmed read data (Ondov et al., 2016). SPAdes assemblies were generated from the high-quality genome sequence data for each isolate (Bankevich et al., 2012), evaluated with QUAST (Gurevich et al., 2013), and the best preexisting genome assembly was identified with the Mash distance from all genomes available in NCBI RefSeq (O’Leary et al., 2016) and GenBank^®^ (Clark et al., 2016) at the time of this study (Supplementary Tables 3 and 4). Multiple genome alignments were performed with progressiveMauve to compare gene gain, loss, and rearrangement of the *E. faecium* isolate (Darling et al. 2010, Supplementary Figure 1). Genes were annotated *de novo* with prokka (Seemann, 2014) from assembled contigs or detected by alignment with ABRicate (Zankari et al., 2012) using a custom database of expected genes based on prior annotations from known strains. Taxonomic analysis of metagenomes and metatranscriptomes was performed by Bowtie2 (Langmead and Salzberg, 2012), SAMtools (Li et al., 2009), and Qualimap2 (Okonechnikov et al., 2016) mapping of reads to the expected genomes, and Mash Screen (Ondov et al., 2019) containment estimations with a custom reference genome database. When equivalent hits were identified for one genome, the larger number was selected to simplify final reporting and analysis. Bowtie2 was used to map reads to a custom database of genes present within the isolates. Metagenomic assembly was performed with metaSPAdes (DNA) (Nurk et al., 2017) or rnaSPAdes (RNA) (Bushmanova et al., 2019) before downstream gene annotation with prokka and ABRicate. Metagenomic reads derived from DNA extracted from contact scenarios and RNA from controlled spike-ins of pathogen to background organisms aligned to genes within the custom database using Bowtie2. Data analysis figures were generated with R Studio, all sequence data were submitted to NCBI BioProject 530203, and intermediate analysis results from the tools can be found at OSF project https://osf.io/3qwps/.

## 5 Results

### 5.1 Modeling Transmission

Bacterial transmission can occur through either a direct skin-to-skin situation, such as between an infected patient and a health care worker, or an indirect skin-to fomite-to skin scenario, where an infected patient touches an object (i.e., cotton, nitrile, stainless steel surface) followed by a healthcare worker touching the same surface. To simulate these two contact scenarios, approaches were developed in Weber et al. 2020 to investigate direct and indirect ESKAPE+C pathogen transfer utilizing a synthetic human skin material VITRO-SKIN^®^ N-19 with or without including a relevant handwashing step. As pathogen transfer does not occur in isolation, a set of background bacteria were included on the VITRO-SKIN^®^ coupon to represent the native skin microbiota that could be present on human hands. Supplementary Tables 1-6 and Supplementary Figures 1-2 describe features of the ESKAPE+C pathogen and background organisms analyzed, as well as their closest matching reference genome in NCBI databases. While culture data was successfully collected from all direct and indirect scenarios previously described (Weber et al., 2020), the indirect wash scenarios were not sequenced in this study because it was anticipated that they would fall well below the sequencing limit of detection (Figure 1).

### 5.2 Taxonomic Detection of Pathogens in Metagenomes

Metagenomic analysis successfully identified the commensal and pathogenic organisms present at the high spike-in level (∼10^6^ CFU/mL), direct contact scenarios. Taxonomic analysis included the use of read mapping and containment estimations with a custom reference genome database that consisted of the specific strains used in this study (Supplementary Tables 3 and 4). As identified in Figure 2, the Mash Screen identity value of 0.90 served as a reasonable threshold for bacterial genome detection in metagenomes of this study. The indirect and low spike-in scenarios did not result in enough sequence coverage of the bacterial genomes to lead to a positive detection event (Figure 3). Ultimately, pathogen abundance within the overall microbial community played the largest role in successful taxonomic classification.

**Figure 2.**
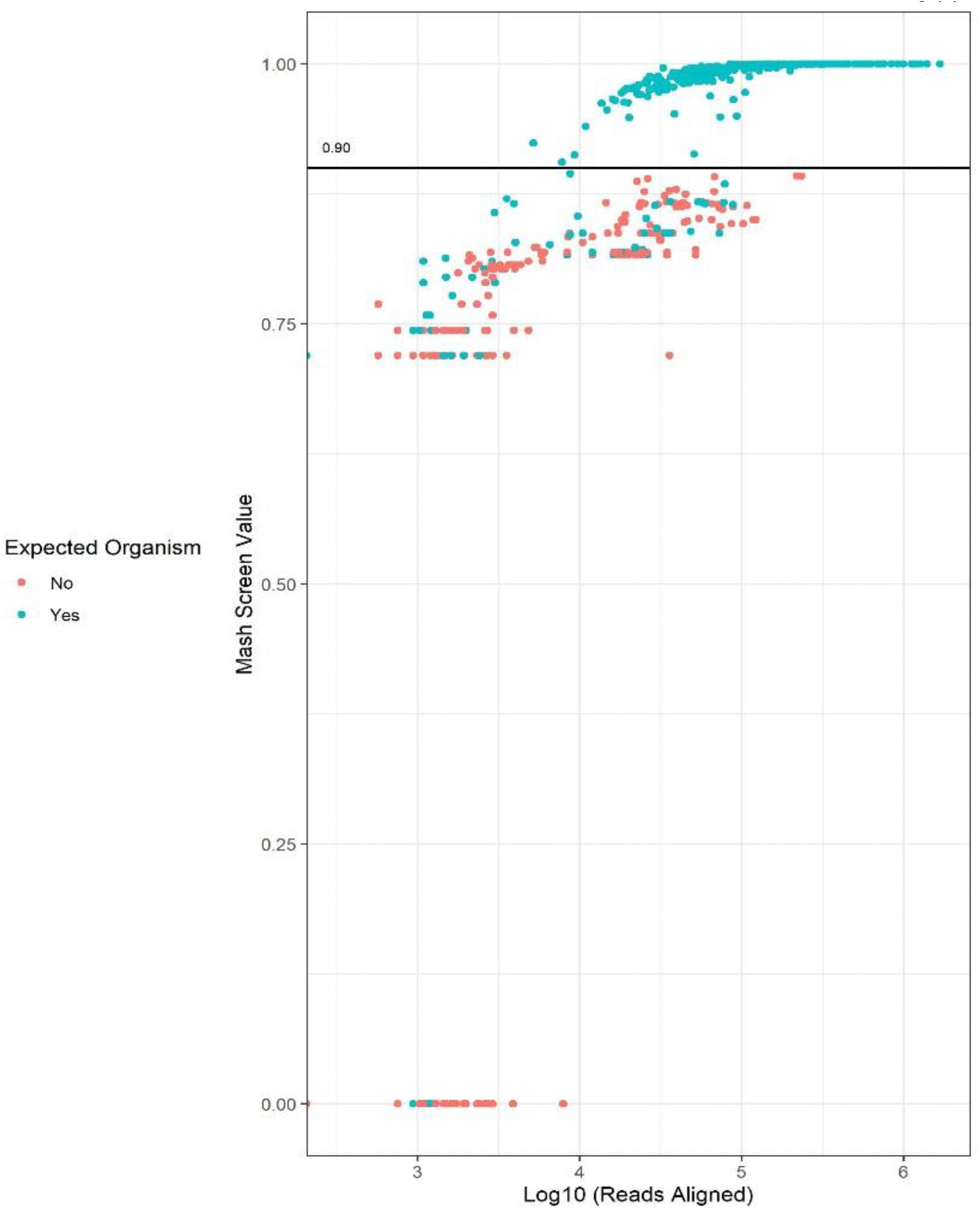
Mash Screen Identity Threshold for Overall Pathogen Detection in Metagenomes. A Mash Screen identity threshold of 0.90 served as a reasonable threshold for bacterial genome detection in metagenomes of this study with a custom database of the expected genome strains. While some true positives (green) fell below the 0.90 threshold, none of the false positives (red) were detected above it. Two or three ESKAPE+C pathogens were cultured together in three different mixes before sequencing, and because the Mash Screen custom database only contained the ESKAPE+C and background organisms, the false positives shown here represent pathogens that were not part of the particular mix that was sequenced.

**Figure 3.**
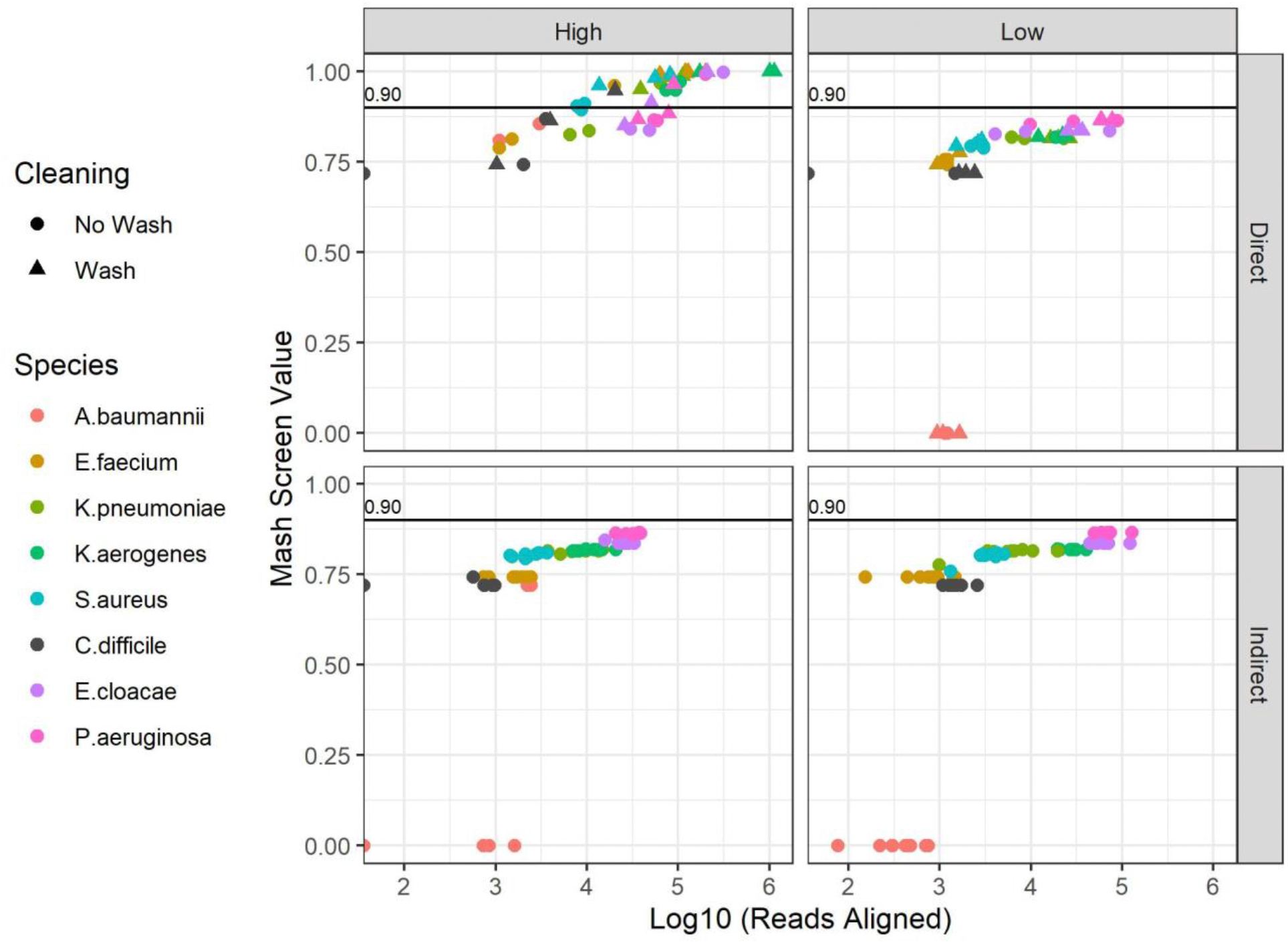
Pathogen Detection in Metagenomes after Different Simulated Contact Scenarios. The Mash Screen identity detection threshold (y axes) of 0.90 (labeled solid black lines) was only met by a portion of the metagenomes sequenced from the direct, high-spike-in scenarios under both wash and no wash conditions. The indirect and low spike-in scenarios did not result in enough coverage of the bacterial genomes to lead to a positive detection event. Alignment of reads to the expected pathogen genomes showed a similar trend of stronger genomic signals in the direct, high-spike-in scenarios (x axes). Notably, *C. difficile* (black) was difficult to detect with metagenomics even in high-spike-in scenarios, which may be due insufficient yield of DNA from its endospore state.

### 5.3 Impact of Simulated Handwashing on DNA Yield

ESKAPE+C pathogens were cultured together in three different mixes based on their media and growth requirements (Weber et al., 2020). Samples were split between the culturing and metagenomics sequencing experiments, and the results were compared downstream. Two or three pathogens were included in each cultured mix, and eight background microorganisms were included along with the pathogens before each metagenome was sequenced (Figure 1). Therefore, the reads within a metagenome that did not map to an ESKAPE+C reference genome in Table 1 and Figure 4 had originated from either 1) another pathogen in the mix or 2) one of the background microorganisms. The CFU/mL values calculated from the culture data were compared to the percentage of metagenomic reads mapped to one of the ESKAPE+C reference genomes. Pathogens were only detected from metagenomics data in the high spike-in direct contact scenarios (Figure 3), such as direct transfer events with and without handwashing. There was no observable correlation between CFU/mL and reads mapped to the reference genomes (Supplementary Figure 3).

**Table 1.**
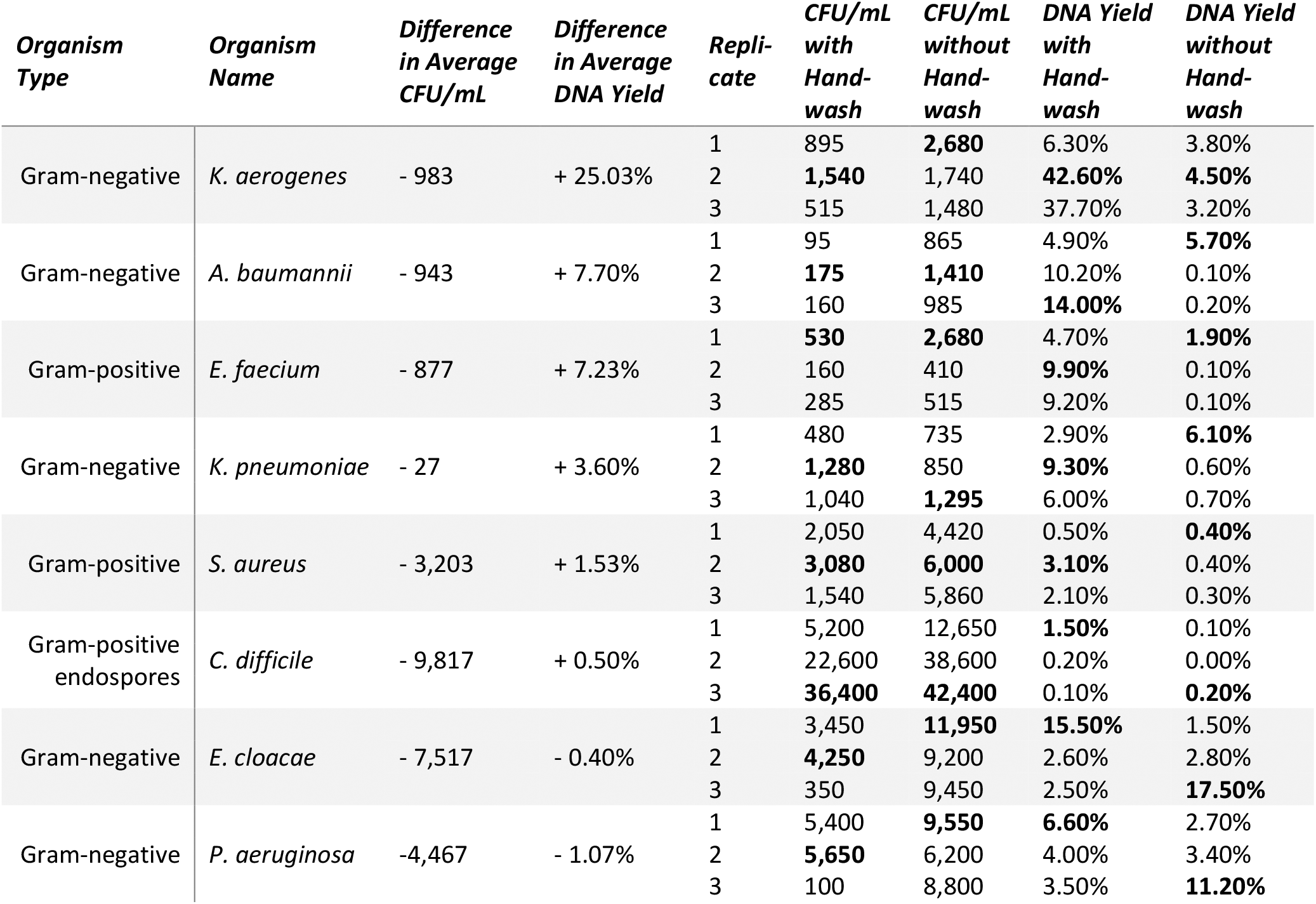
Effect of Handwashing on ESKAPE+C Cultures and DNA Sequence Recovery. The data points in Table 1 were collected in three replicates from the direct contact, high spike-in scenarios. Note that although the three replicates for handwash and no handwash events are listed on the same rows in Table 1, they were independent events. The highest values for each organism per column are bolded. DNA yield refers to the percentage of reads in a metagenomic sample that aligned to the reference pathogen genome. The difference in average DNA yields between handwashing and no handwashing replicates is listed in the third column, subtracting the average DNA yield from the no handwash replicates from the average DNA yield in the handwash replicates for each pathogen. Positive values indicate the average DNA yield was higher in handwash than no handwash, and negative values indicate average DNA yields were higher in no handwash scenarios. Similarly, average CFU/mL differences are listed for handwash and no handwash scenarios.

**Figure 4.**
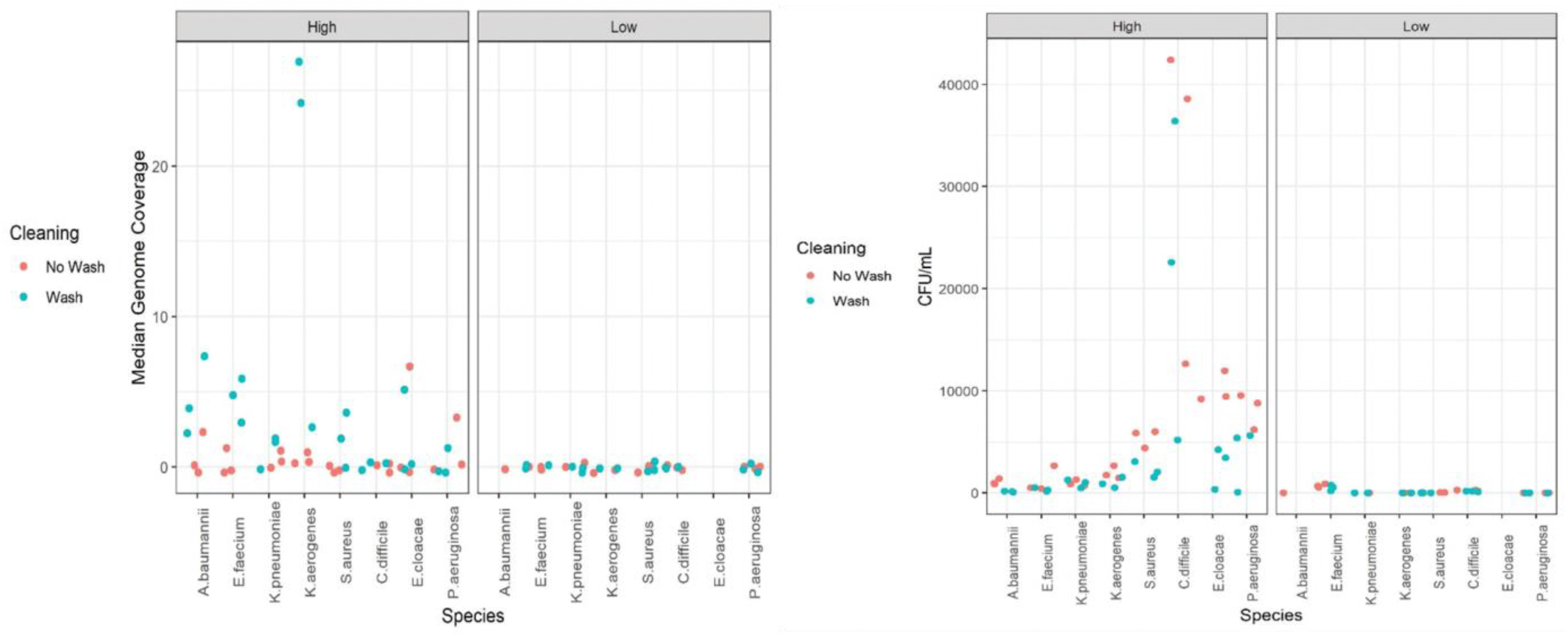
Influence of Handwash on DNA Abundance and ESKAPE+C Viability. Looking at all of the culturing and metagenome sequence data together, handwash (blue) tended to have an opposite effect on median genome coverage – increasing the amount of ESAKPE+C DNA yielded for sequencing (left panels) and decreasing viable the number of viable ESAKPE+C pathogens detected as CFU/mL in culture data (right panels). Conversely, no handwash (red) decreased DNA yield from the pathogens and resulted in more viable pathogen CFUs. This trend was only visible in the high spike-in scenarios (∼10^6^ CFU per 9 cm^2^ coupon), as pathogens spiked in at low amounts (≤10^4^ CFU per 9 cm^2^ coupon) fell below the limit of detection with metagenomic sequencing. Culture data still picked up some signal in these low spike-in instances (1-904 pathogen CFU/mL after transfer for direct contact scenarios and 1-72 pathogen CFU/mL for indirect contact scenarios). Metagenomics detected pathogens for only the high spike-in direct no wash scenarios (average of ∼2,447 pathogen CFU/mL after transfer), while culture data detected pathogens in all high spike-in scenarios (95-42,400 pathogen CFU/mL after transfer for direct contact scenarios and 1-1,600 pathogen CFU/mL for indirect contact scenarios).

While the simulated handwashing events on VITRO-SKIN^®^ decreased the number of viable pathogen cells, they resulted in higher overall pathogen detection relative to the non-handwashing scenarios rates within metagenomes in the high spike-in (∼10^6^ CFU/mL) direct contact scenarios (Table 1, Figure 4). Most ESKAPE+C species yielded more DNA after the handwash step compared to no handwashing, suggesting that the handwash helped to lyse the bacterial cells and release more DNA for sequencing. Gram-negative *Klebsiella aerogenes* (formerly known as *Enterobacter aerogenes*), *A. baumannii*, and *K. pneumoniae*, as well Gram-positive *E. faecium*, showed the largest maximum increases in DNA yield after the simulated handwashing scenarios. Gram-positive *S. aureus* and *C. difficile* endospores also showed a relatively smaller impact of handwashing on increased DNA yield. Gram-negative *P. aeruginosa* and *Enterobacter cloacae* were the exceptions, with no handwash scenarios showing a larger maximum or average DNA yields compared to handwash scenarios. The relatively thin layers of peptidoglycan in the Gram-negative cell walls of *P. aeruginosa* and *E. cloacae* may have allowed the microorganisms to be more effectively lysed and DNA recovered with or without the handwash step, although this trend was not observed for all Gram-negative bacteria in Table 1.

### 5.4 Detection of Antimicrobial Resistance and Virulence Genes in Metagenomes

To evaluate rates of gene detection and coverage, metagenomic reads derived from contact scenarios were mapped to the custom database of genes that were annotated within the assembled genomes of ESKAPE+C pathogens and background organisms (Figure 5, Supplementary Figures 4-7). Like the ESKAPE+C genome-level analyses, there was a decrease in pathogen gene signals in the indirect contact scenarios compared to the direct contact scenarios (Figure 6). The wash with high inoculum detected more reads from genes compared to no wash scenarios, and there was no detection of pathogen genes in low inoculum. Though similar to the genome-level analyses, *E. cloacae* was an exception to this in that no handwash scenarios showing a larger maximum DNA yield compared to handwash scenarios. Antimicrobial resistance (AMR) genes detected within the contact scenarios included several that encode proteins specific for antibiotic inactivation including aminoglycoside resistance genes *aac(6’), -ach(2”)*, and *aadD* from *S. aureus, aac(6’)-li* and *ant(6)-Ia* from *E. faecium*, the macrolide resistance gene *ere(A)* from *E. cloacae*, and the lincosamide resistance gene *lnuB* from *E. faecium*. The *qnrS1* gene encoding an antibiotic target protection protein *cloacae*.

**Figure 5.**
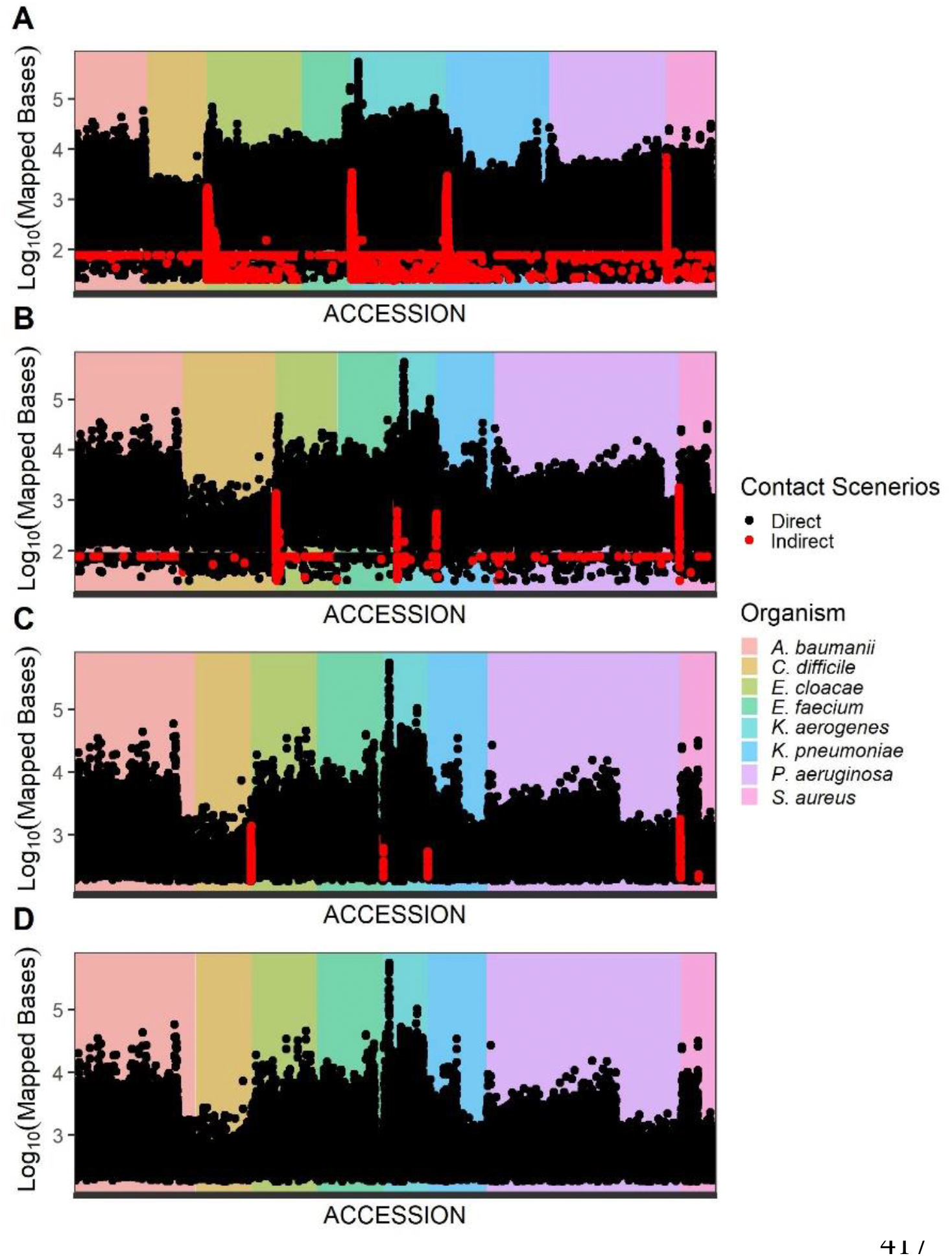
Comparing DNA Reads Mapped to Genes in Direct and Indirect Contact Scenarios. Sequences from direct contact (black points) and indirect contact (red points) scenarios were mapped to nucleotide sequences of genes in pathogens within the custom database. This figure is labeled according to (A) all mapped reads, (B) removal of shared genes (cross-hits to each other), (C) application of a minimum of 180 bp mapped bases cutoff, and (D) removal of 52 accessions considered false positives (accession hits associated with a different pathogen mixture). Even before removal of accessions related to known false positives and shared genes, there was a significant decrease in gene signals related to the pathogens in the indirect contact scenarios compared to the direct contact scenarios. This could be attributed to the overall significant decrease in the amount of pathogen DNA present after transfer events in the indirect contact scenarios.

**Figure 6.**
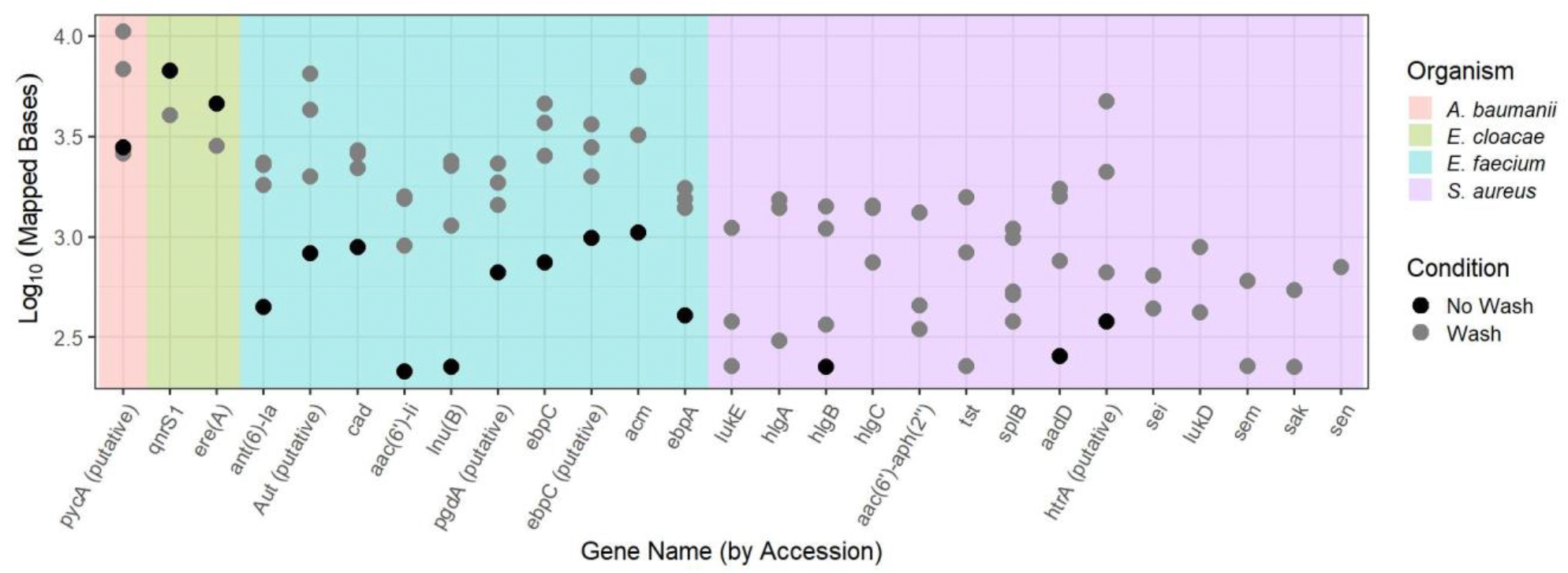
DNA Reads Mapped to a Selection of Potential Virulence and AMR Genes in Direct Handwash and No Handwash Scenarios. Metagenomic sequences from direct transfer scenarios were mapped to genes related to AMR and virulence. The majority of detected AMR and virulence genes from contact scenarios were from *E. faecium* and *S. aureus*, while *E. cloacae* and *A. baumannii* were also detected in their respective mixtures. Like the genome-level analyses, the wash with high inoculum overall detected greater mapped reads compared to no wash and no detection in low inoculum. Also like the genome-level analyses, *E. cloacae* was an exception where no handwash scenarios showed a larger maximum DNA yield compared to handwash scenarios. Gram-positive *S. aureus* had a relatively smaller impact of handwashing on increased DNA yield when read mapping at the genome-level (Table 1), but the impact of handwashing appears relatively larger when viewed at the AMR gene level. Although completely shared genes (100% length and identity) across multiple species were removed from analysis, it is possible that partially shared genes accounted for the increased number of mapped short reads at the AMR gene level.

Genes encoding virulence outnumbered those encoding AMR. For *S. aureus*, encoded cytolytic factors included the γ-hemolysin encoding operon *hlgABC* and the *lukDE* genes encoding leukotoxins (Dumont et al., 2011; Vandenesch et al., 2012; Alonzo et al., 2013). Other genes that encode damaging virulence factors include *tsst*, the inflammation-inducing toxic shock syndrome protein (Brosnahan and Schlievert, 2011; Kulhankova et al., 2018), and the enterotoxin/superantigen-encoding genes *sei, sem*, and *sen* (Omoe et al., 2013; Roetzer et al., 2016; Ono et al., 2017). The *sak* gene codes for the immune-subverting anti-host complement, anti-immunoglobulin, and *anti*-antimicrobial peptide effector staphylokinase (Rooijakkers et al., 2005b, 2005a; Kwiecinski et al., 2013). Virulence genes detected in *E. faecium* included the collagen adhesin gene *acm* (Nallapareddy et al., 2003), the *ebpA* and *ebpC* genes for endocarditis-and-biofilm-associated pilus genes (Nallapareddy and Murray, 2008; Nallapareddy et al., 2011; Montealegre et al., 2015), and virulence response genes encoding peptidoglycan N-acetylglucosamine deacetylase A encoded by *pgdA* (Benachour et al., 2012). ABRicate detection of gene coverage in assembled contigs demonstrated similar patterns of pathogen-specific differences in the loss of signal between high inoculum handwash and no handwash events (Figure 7). The detection of signal from *C. difficile* with no wash was negligible, while the signal from *P. aeruginosa* and *A. baumannii* were successfully retained in no wash events. Like the genome mapping results, *P. aeruginosa* showed a stronger signal after no handwash than handwash, which was different than the majority of the other ESKAPE+C pathogens that increased in DNA yield after the simulated handwashing events. The greater detection and difference in loss of signal from different organisms suggest that metagenomic methods are significantly affected by the physicochemical features of the organisms, and the circumstances employed before DNA extraction.

**Figure 7.**
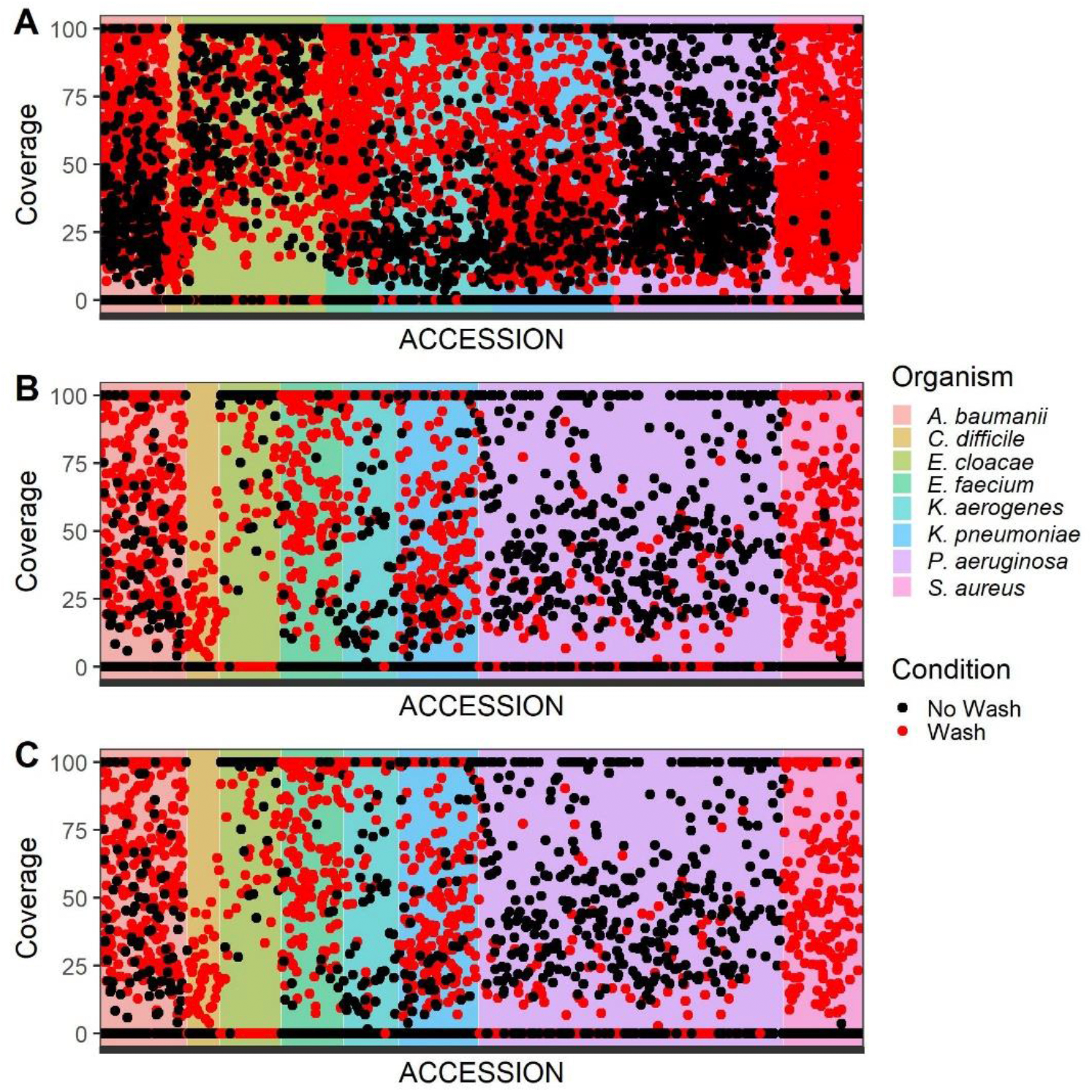
ABRicate Coverage of ESKAPE+C Genes Identified within Assembled Metagenomes in Direct Handwash and No Handwash Scenarios. This figure is labeled according to (A) all genes identified by ABRicate within the assembled contigs using the custom gene database, (B) after removal of shared genes, (C) after removal of 52 accessions known as false positives. Remaining accessions included direct contact scenarios of high inoculum that were simulated with and without handwash. ABRicate detection of gene coverage demonstrates pathogen-specific differences in the loss of signal between handwash and no wash events.

### Detection of ESKAPE+C Pathogens in Titrated Metatranscriptome Experiment

Mapping of metatranscriptomic reads derived from RNA extracted from different concentrations of ESKAPE+C pathogens spiked into a constant concentration of background organisms demonstrated a loss of signal as ESKAPE+C bacterial concentration decreased (Figure 8). The relationship of mapped bases and mean coverage in relation to estimated CFU from the spike-ins of ESKAPE+C pathogens to background organisms was dependent on the organism. As a general trend, gene signals were detected only for bacteria spiked in at the highest level (∼10^6^ CFU/mL) (Figure 9). Only seven of the AMR and virulence genes that were detected in the contact scenarios were detected in the extracted RNA from pathogen spike-ins (Supplementary Figure 8), suggesting low expression of these genes. Low gene expression is not unexpected, given the absence of selection pressure and recent contact with a host organism. Different genes specific to a microorganism demonstrated variability, while some within the same microorganism showed a positive direct linear relationship with the CFU spike-in level. Other genes within a microorganism appeared consistent as estimated CFU increased (Supplementary Figure 9). Interestingly, metatranscriptomic analysis of the pathogen spike-in detected specific genes from *C. difficile*, while the *C. difficile* signal was absent for genome detection methods (Figure 10). The detection of mRNA related to sporulation (i.e., small acid-soluble protein, spore coat protein) in *C. difficile* within spike-ins indicates the potential for determination of the presence of specific gene transcripts that can be contributed to a genus or species within a mixture depending on the gene or partial sequence specificity to a taxonomic level.

**Figure 8.**
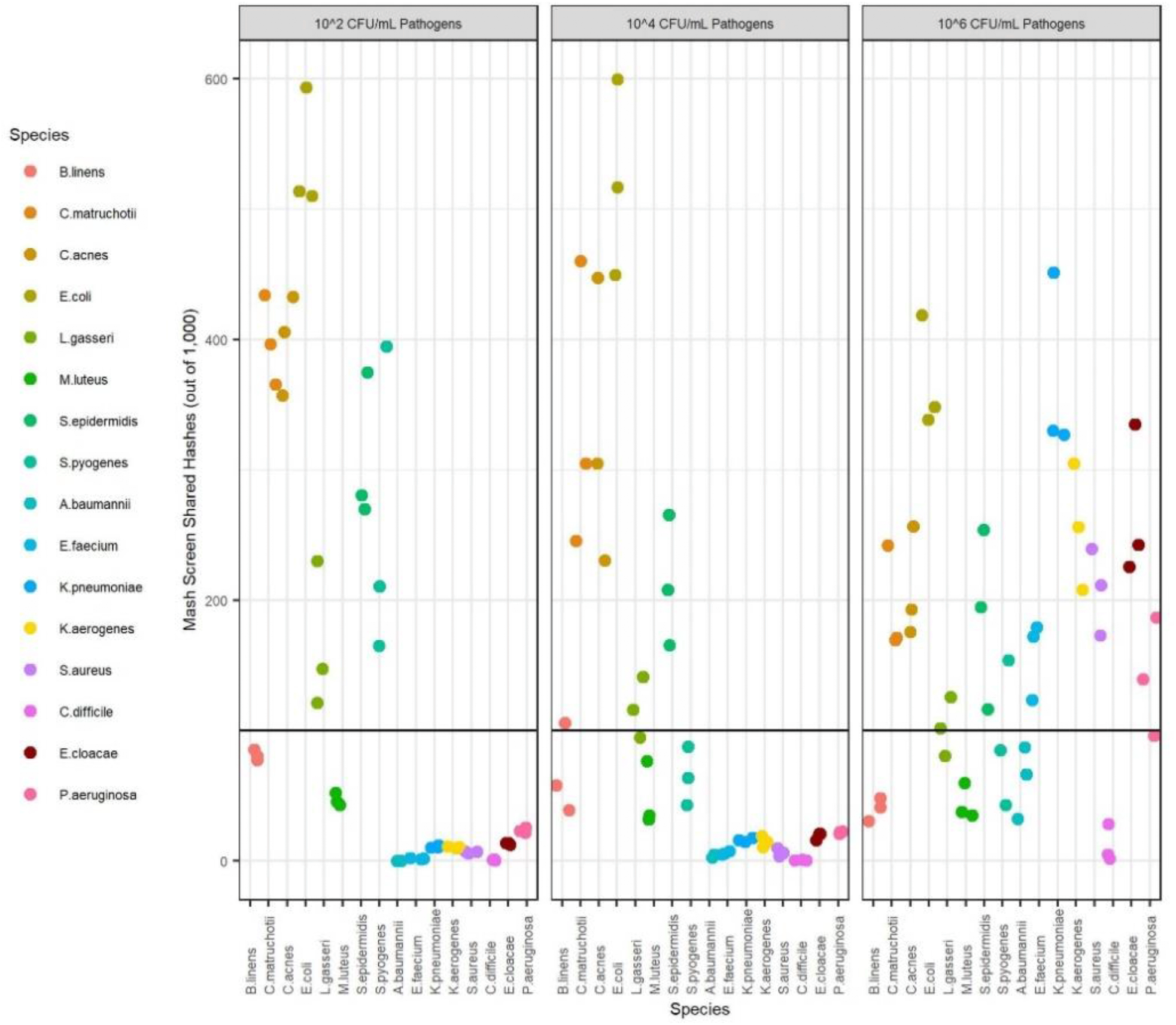
ESKAPE+C Pathogen Detection Limits in Metatranscriptomes. A horizontal black line is drawn to show the equivalent of a successful genome-level detection event at a Mash Screen identity of 0.90. ESKAPE+C pathogens within metatranscriptomes were not detected below 10^6^ CFU/mL in the RNA titration experiments. Only the background microorganisms were detected, which were kept at a constant level of 10^6^ CFU/mL in all samples. *C. difficile* and *A. baumannii* were not detected in metatranscriptomes at any tested level.

**Figure 9.**
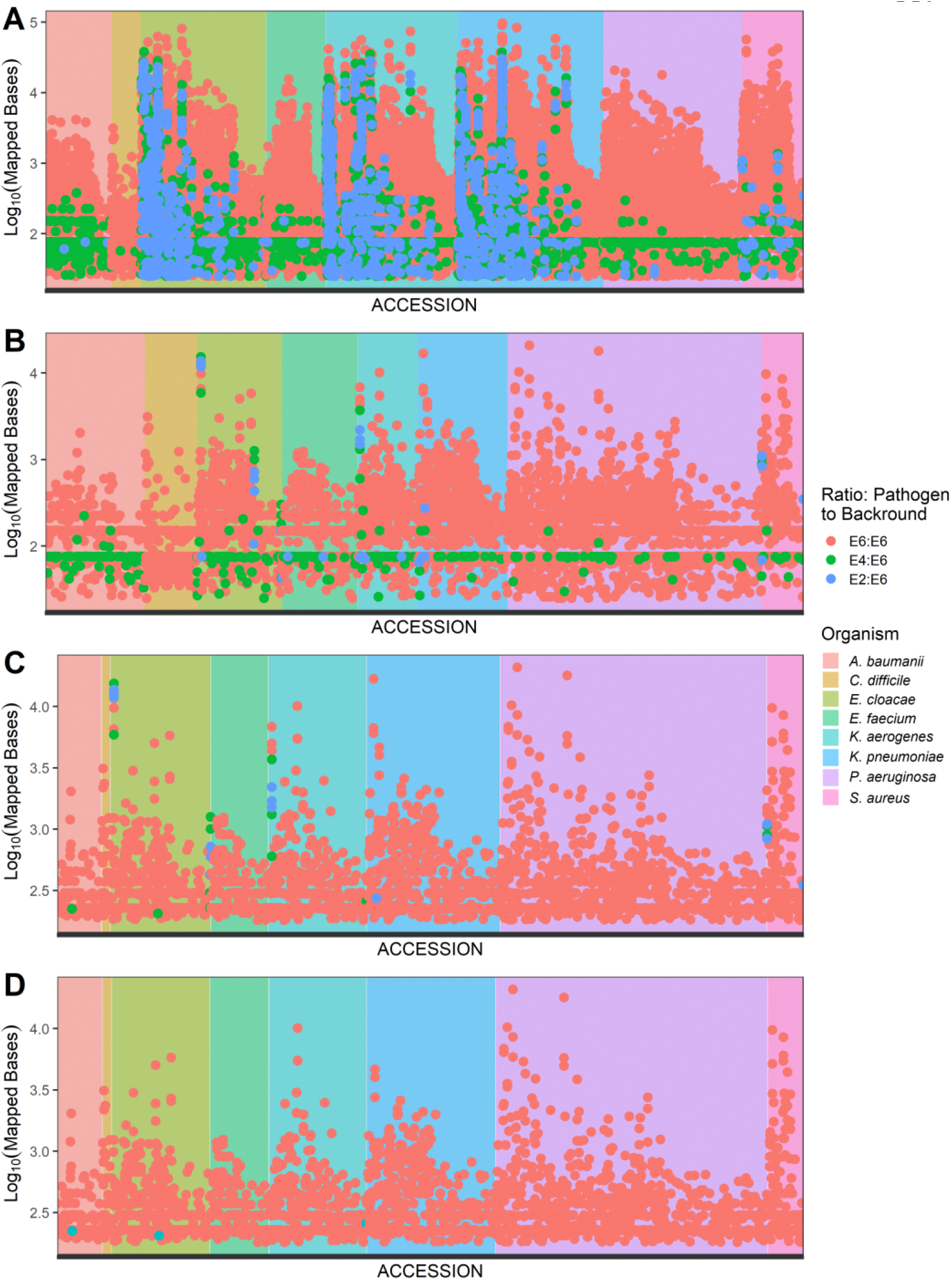
ESKAPE+C Pathogen RNA Reads Mapped to Genes. Reads were mapped from RNA titration experiments to nucleotide sequences of pathogen genes within the custom database. This figure is labeled according to (A) all mapped reads, (B) removal of shared genes, (C) application of a minimum of 180 bp mapped bases cutoff, and (D) removal of 52 accessions considered false positives, as described previously. This clearly shows the importance of considering shared gene content before attributing the origin of a gene to a specific species, as many of the low spike-in level genes (blue) in panel A disappeared in panel B. After all quality control filtering steps were completed, the genes identified were only those from the high spike-in level in panel D.

**Figure 10.**
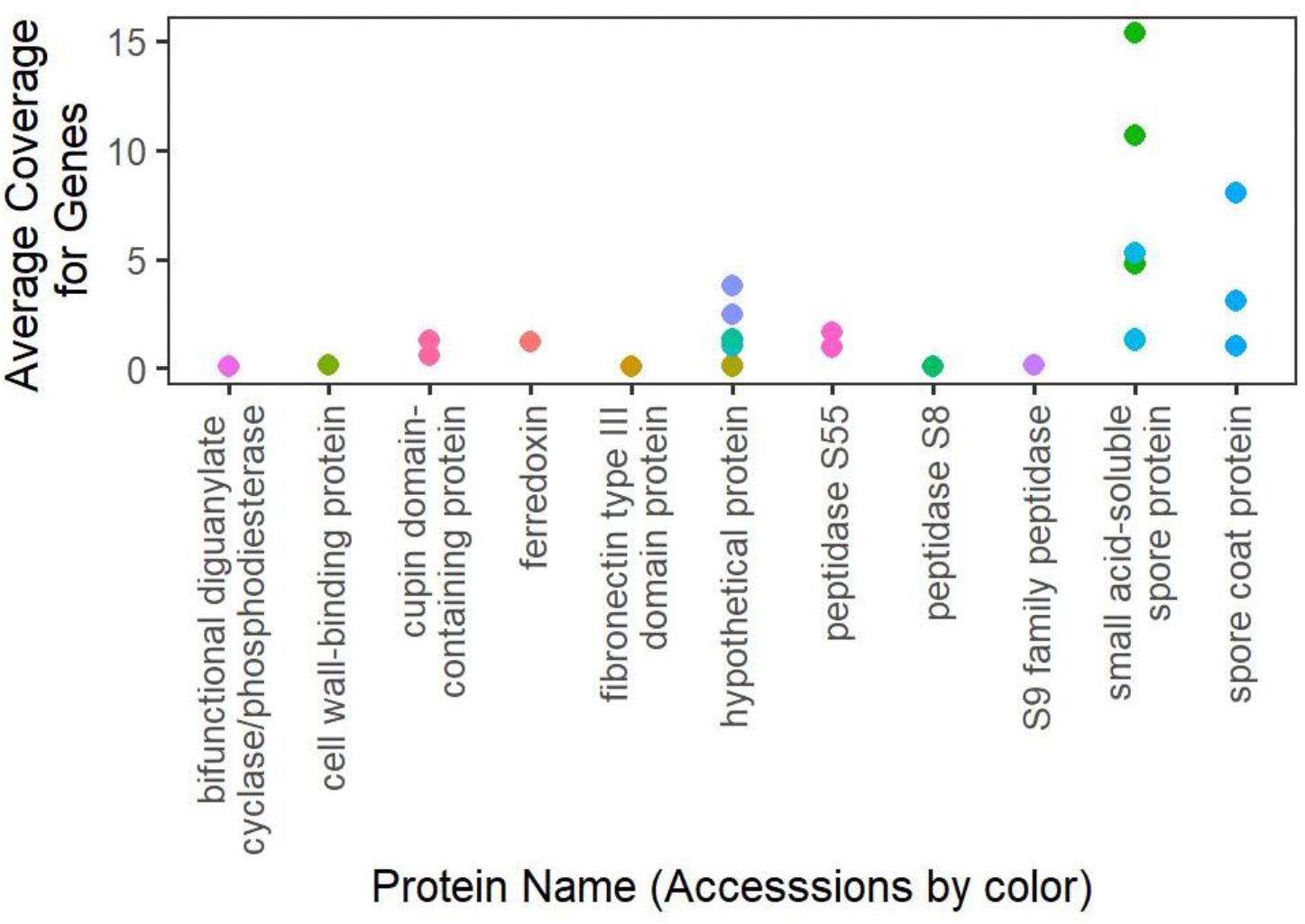
RNA Coverage of *C. difficile* Genes. RNA reads were mapped to *C. difficile* genes within the custom database. Genes for *C. difficile* were only detected at the highest spike-in level of 10^6^ CFU/mL. Different gene accessions for the same gene descriptions are depicted in different colors. The detection of mRNA related to sporulation in *C. difficile* indicates the potential for determination of the presence of specific genes transcripts that could be contributed to a genus or species within a mixture, depending on the specificity of a gene or partial sequence to a taxonomical level. It also makes sense that sporulation genes would be expressed, since *C. difficile* endospores were used in the experiments.

## Discussion

Culturing of bacteria is commonly performed to identify infectious pathogens (Nekkab et al., 2017), but culture-dependent methods have inherent limitations. Established nucleic acid-based detection approaches like polymerase chain reaction (PCR) or 16S rRNA sequencing overcome some of these limitations, including the requirement for pathogen viability or culturability; however, these targeted methods are only able to detect limited, known genomic regions and fall short of identifying pathogens at the strain or even species level. Metagenomic analysis methods hold substantial promise in overcoming these hurdles. By drawing upon an established, well-characterized data set, we have illustrated many of the strengths and weaknesses posed by unbiased metagenomic and metatranscriptomic sequencing. Although unbiased sequencing methods have enormous potential for pathogen detection, this study has highlighted limitations in failing to detect low abundance pathogens in mixed samples (≤ 10^4^ CFU/mL), challenges with determining the species of origin for gene sequences that are partially or fully shared among multiple species (e.g., AMR genes), and variable rates of nucleic acid extraction efficiency in different bacteria.

After mapped sequences of DNA extracted from different contact scenarios were compared to previously reported ESKAPE+C colony counts through selective culturing, the mapped reads in direct transfer scenarios demonstrated considerable variation in which pathogen species were detected and how the conditions of simulated handwash or no handwash impacted the results. Only three replicates were performed per condition in this study, and it is possible that additional technical replicates could reduce the amount of observed variability. However, additional technical replicates would not overcome variation due to inherent biological characteristics, like differences in cell wall and membrane structures influencing the amount of DNA extracted (e.g., *C. difficile* endospores). Differences in NGS library preparation methods can also introduce bias in the detection of different bacterial species (Morgan et al., 2010; Van Dijk et al., 2014), and the results from handwash vs. non-handwash scenarios may be conceptually similar to different sample processing and cell lysing methods before sequencing. Bias can also be introduced in CFU measurements, as selective media utilize intrinsic attributes of a bacteria to isolate and differentiate from other bacteria, and this selective pressure may also eliminate viable or injured bacteria that are unable to recover once plated (Apajalahti et al., 2003).

It was promising that genes were detected within microbial mixtures despite the lack of detection with culturing techniques or genome-level taxonomic calls. This suggests that functionally informative genes (e.g., virulence, AMR) could have lower the limits of detection than culturing or standard taxonomic identification methods. *A. baumannii* was at or near the limit of detection for the culturing method (left of dotted line in Supplementary Figure 5), while specific genes from these species were detected by sequencing. Compared to whole genome techniques, the method of assembly and annotations of organism specific genes led to greater retention of pathogen signal, especially from the no handwash scenarios. The comparison of ABRicate annotations of assembled contiguous sequences with Bowtie2-mapped bases demonstrates that while alignment of short reads allowed for more sensitive detection of genes, the annotation of assembled reads provided better precision in gene identifications. High precision was especially true when at or near 100% coverage was achieved for a species-specific gene within a contiguous sequence.

Bacterial nucleic acids from skin and nonsterile specimens may result in a stronger signal than that of the pathogen (Gu et al., 2019), and in this study that was simulated by a constant level of “background” bacterial species. The lack of signal from ESKAPE+C pathogen sequences within the indirect scenarios using unbiased NGS methods compared to selective culturing techniques could be attributed to the lack of selection for pathogen signal over background microorganism signal in greater quantity, highlighting a key challenge of NGS in clinical settings with nonsterile specimens and complex sample types. Enrichment techniques to amplify the pathogen signal before sequencing could have improved the limits of detection for ESKAPE+C in all scenarios, but enrichment techniques often come at the price of biased sequencing in search of known targets. Such methods have limited applicability to emerging or novel pathogens, as well as situations when the infectious agent is unknown to the physician and fails standard clinical tests.

## 6 Conclusions

Metagenomic and metatranscriptomic analyses promise an unbiased approach to pathogen species-level detection and functional gene characterization within clinical samples; however, the limitations of this technology must be fully evaluated before traditional culturing methods can be supplemented or replaced by this new methodology. This study makes significant progress toward this goal, capitalizing on a large, well-curated data set from a previously published study and generating complimentary NGS analyses for comparison. In doing so, we illustrate both the strengths of this type of analysis, such as the ability to identify pathogens and characterize elements of virulence or the resistome in a given sample, as well as the limitations of unbiased sequencing, predominantly highlighted by low sensitivity when pathogens are present at low abundance within a complex mixture. Variability in the loss of signal from different bacterial species also lends support to how laboratory and bioinformatics methods are impacted by the intrinsic nature of the organisms, as it relates to nucleic acid extraction and the uniqueness of genome content. These results will inform and aid the healthcare and epidemiological community as they evaluate the appropriate scenarios to utilize metagenomic analysis.

## 7 Abbreviations

HAI: Healthcare-acquired infections
NGS: Next Generation Sequencing
ESKAPE+C: *Enterococcus faecium, Staphylococcus aureus, Klebsiella pneumoniae, Acinetobacter baumannii, Pseudomonas aeruginosa, Enterobacter* species and *Clostridioides difficile*
CDC: Centers for Disease Control and Prevention
AR: Antibiotic resistance
AMR: Antimicrobial resistance
ATCC: American Type Culture Collection
CFU: Colony forming units
PBS: Phosphate buffered saline
mRNA: messenger ribonucleic acid
mL: milliliter
polymerase chain reaction: PCR
*Enterococcus faecium*: *E. faecium*
*Enterobacter cloacae*: *E. cloacae*
*Staphylococcus aureus*: *S. aureus*
*Klebsiella pneumoniae*: *K. pneumoniae*
*Klebsiella aerogenes*: *K. aerogenes*
*Acinetobacter baumannii*: *A. baumannii*
*Pseudomonas aeruginosa*: *P. aeruginosa*
*Clostridioides difficile*: *C. difficile*
*Brevibacterium linens*: *B. linens*
*Corynebacterium matruchotii*: *C. matruchotii*
*Cutibacterium acnes*: *C. acnes*
*Escherichia coli*: *E. coli*
*Lactobacillus gasseri*: *L. gasseri*
*Micrococcus luteus*: *M. luteus*
*Staphylococcus epidermidis*: *S. epidermidis*
*Streptococcus pyogenes*: *S. pyogenes*

## 8 Acknowledgements

The authors would like to thank Mr. Jim Gibson for his assistance creating the contact scenario figure and Dr. Madeline Roman for her review of this manuscript. The authors would also like to thank Drs. Alison Laufer Halpin and Rachel Slayton of the CDC for their constructive feedback throughout this project.

## 9 Author Contribution Statement

All authors have read and approved the manuscript. Study Conceptualization, KLT, NCK, DSL, KLW, FCH; Laboratory Analysis, KLW, DSL, KQS; Bioinformatics, KLT, NCK, VP, GDG, ADK, CAA; Project Administration, KLT, FCH; Writing (Original Draft Preparation) and Visualization, FCH, KLT, ADK, NCK, DSL, KLW; Writing (Review & Editing), KLT, FCH, GDG, DSL, ADK, GDG.

## 11 Availability of data and materials

All data generated or analyzed during this study are included in this published article, its supplementary files, our OSF project (https://osf.io/3qwps/), or the NCBI BioProject 530203.

