## Supplementary Information for "Detection of ESKAPE pathogens and *Clostridioides difficile* in Simulated Skin Transmission Events with Metagenomic and Metatranscriptomic Sequencing"

**Supplementary Table 1: Pathogen Isolates**

| Species | Pathogen Source | Sequenced In-House? | Genome Coverage | NCBI Sequence Identifier | Laboratory Mix* |
|---|---|---|---|---|---|
| *Acinetobacter baumannii* | CDC AR Bank #0275 | No | 63x | SRR4417583 | Mix 1 |
| *Enterococcus faecium* | CDC AR Bank #0579 | Yes | 81x | SRR10175739 | Mix 1 |
| *Klebsiella pneumoniae* | CDC AR Bank #0139 | No | 34x | SRR4025991 | Mix 1 |
| | | | 34x | SRR5168485 | |
| | | | 34x | SRR5168486 | |
| *Klebsiella aerogenes* | CDC AR Bank #0161 | No | 45x | SRR3112300 | Mix 2 |
| *Staphylococcus aureus* | CDC AR Bank #0219 | No | 352x | SRR6985639 | Mix 2 |
| | | No | 44x | SRR4417447 | |
| *Clostridioides difficile*** | ATCC 43598 | Yes | 53x | SRR10175728 | Mix 3 |
| *Enterobacter cloacae* | CDC AR Bank #0365 | No | 132x | SRR6807647 | Mix 3 |
| *Pseudomonas aeruginosa* | CDC AR Bank #0230 | No | 142x | SRR6807660 | Mix 3 |
| | | | 21x | SRR4417531 | |

*Pathogens were cultured together in three different mixes to differentiate and count colonies from each species.
***Clostridioides difficile* was sequenced as endospores throughout this project.

**Supplementary Table 2: Background Organism Isolates**

| Species | Source | Sequenced In-House? | Genome Coverage | NCBI Sequence Identifier |
|---|---|---|---|---|
| *Brevibacterium linens* | ATCC 9172 | Yes | 43x | SRR10175828 |
| *Corynebacterium matruchotii* | ATCC 14265 | Yes | 84x | SRR10175827 |
| *Cutibacterium acnes* | ATCC 11827 | Yes | 59x | SRR10175805 |
| *Escherichia coli* | ATCC 9637 | Yes | 32x | SRR10175794 |
| *Lactobacillus gasseri* | ATCC 33323 | Yes | 122x | SRR10175783 |
| *Micrococcus luteus* | ATCC 4698 | Yes | 34x | SRR10175772 |
| *Staphylococcus epidermidis* | ATCC 12228 | Yes | 61x | SRR10175761 |
| *Streptococcus pyogenes* | ATCC 19615 | Yes | 160x | SRR10175750 |

**Supplementary Table 3: Pathogen Isolate Best Match to a Preexisting Genome Assembly**

| Species | Best Match to a Preexisting Genome Assembly | Strain Name in Preexisting Assembly | Percent Similarity* | Plasmids? |
|---|---|---|---|---|
| *Acinetobacter baumannii* | ASM295131v1 | *Acinetobacter baumannii* Strain R7 | 98.7% | Unknown |
| *Enterococcus faecium* | E8927_hybrid_assembly | *Enterococcus faecium* Isolate E8927 | 88.2% | Yes (6) |
| *Klebsiella pneumoniae* | ASM220219v1 | *Klebsiella pneumoniae* Strain AR_0139 | 96.7% | Yes (4) |
| *Klebsiella aerogenes* | ASM307128v1 | *Klebsiella aerogenes* Strain AR_0161 | 98.2% | Yes (1) |
| *Staphylococcus aureus* | ASM319370v1 | *Staphylococcus aureus* Strain AR_0219 | 98.2% | Yes (1) |
| *Clostridioides difficile* | GCF_900243225 | *Clostridioides difficile* Strain 1470 | 98.7% | No |

| | | | | |
|---|---|---|---|---|
| *Enterobacter cloacae* | ASM296845v1 | *Enterobacter hormaechei subsp. hoffmannii* Strain AR_0365 | 98.5% | Yes (4) |
| *Pseudomonas aeruginosa* | ASM296869v1 | *Pseudomonas aeruginosa* Strain AR_0230 | 97.2% | Yes (2) |

*The percent similarity metric is the “genome fraction” reported by the QUAST -R after aligning our SPAdes assembly to the best preexisting reference assembly.

Although NCBI labels the CDC AR Isolate Bank #0365 isolate “*Enterobacter hormaechei subsp. hoffmannii,”* the CDC AR Isolate Bank labels that species as “*Enterobacter cloacae*.” This seems to be a consistent labeling method that is used by NCBI, since other sequences identified elsewhere as *Enterobacter cloacae* are also labeled as *Enterobacter hormaechei subsp. hoffmannii* within the NCBI framework. Although the best preexisting assembly did not have the expected species name, this assembly was clearly derived from CDC AR Isolate Bank #0365 based on its metadata, and it showed >98% similarity to our SPAdes assembly for *Enterobacter cloacae*. For simplicity and consistency with CDC AR Isolate Bank nomenclature, we will refer to this isolate as *Enterobacter cloacae* throughout this report and the associated project data.

The best preexisting reference assemblies for *Klebsiella pneumoniae*, *Klebsiella aerogenes*, *Staphylococcus aureus*, and *Pseudomonas aeruginosa* have the exact strain names matching to those in the CDC AR Isolate Bank. The strain for *Clostridioides difficile* matches to the expected name within the ATCC documentation (https://www.atcc.org/products/all/43598.aspx). The best preexisting assembly match for *Acinetobacter baumannii* is listed as Strain R7, which is different than the expected name of CDC AR Bank #0275. An NCBI BioSample exists for CDC AR Bank #0275 (SAMN04901665), but it does not have a corresponding reference assembly, which explains why a preexisting assembly match was not found with the exact strain name. We have submitted our assembly for these *Acinetobacter baumannii* CDC AR Bank #0275 reads to NCBI to fill this gap. We have also submitted our assemblies for *Clostridioides difficile* and *Enterococcus faecium* to correspond with our raw reads from sequencing these pathogen strains, particularly since sequence data for this *Enterococcus faecium* strain was not available online prior to this study. The *Enterococcus faecium* Isolate E8927_hybrid_assembly was the best preexisting assembly match for CDC AR Isolate Bank #0579, but it had only 88.2% similarity (i.e., QUAST -R genome fraction) to the SPAdes assembly from our high-quality isolate sequences. To further explore this low genome fraction percentage, and to examine the overall genomic plasticity of E. faecium, 120 NCBI *Enterococcus faecium* complete assemblies were aligned to each other using Mauve. Mauve allows for visualizations of genome-wide evolutionary dynamics, including rearrangements and inversions (Supplementary Figure 1).

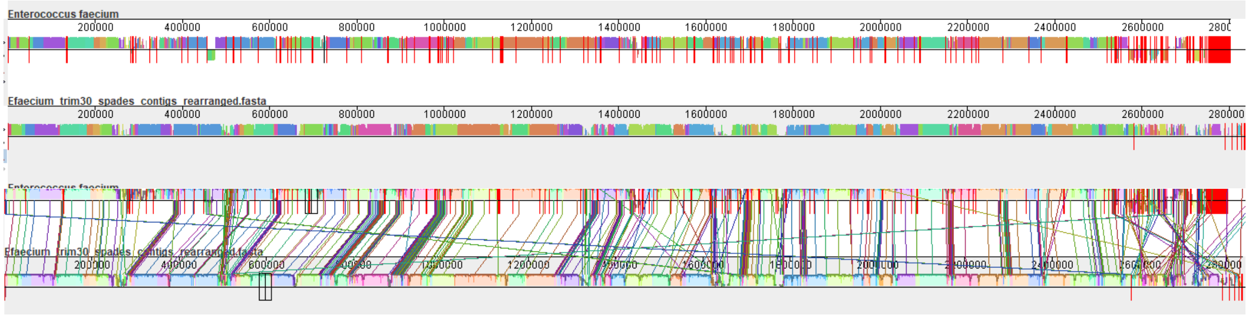

**Supplementary Figure 1. Mauve Visualization of *Enterococcus faecium* Genome Assembly Alignments.** Visualization of genome alignments between our CDC AR Bank #0579 SPAdes assembly (top genome) and the preexisting E8927 assembly (bottom genome). Alignments are shown without lines (top image) and with lines (bottom image), which connect regions of locally collinear blocks with sequence similarity. Red vertical lines represent breaks between contigs for the SPAdes assembly and breaks between chromosome and plasmids for the preexisting reference assembly.

The multi-genome alignments for 120 *Enterococcus faecium* complete assemblies confirmed that E8927 was the best preexisting reference assembly for CDC AR Isolate Bank #0365. The E8927 assembly was similar in length, as well as in the order of sequence blocks, to the other available *Enterococcus faecium* complete assemblies. There was variability in the sequence similarity in single or operonic genes within the individual sequence blocks, which could be due to insertion or deletion of genes and operons. The highest variable region of the complete genomes was the end of the assemblies, where plasmids would be represented, if present. The E8927 assembly has six plasmids, and since this relatively large number of plasmids may vary from assembly to assembly, this could be a significant source of sequence misalignments between assemblies. The alignment of all 120 assemblies also showed evidence of a potential split within *Enterococcus faecium*, represented by a major inversion event of approximated 1.4 Mb. It is possible that the characteristic multi-clade structure of the species may be attributed to this large inversion event.

**Supplementary Table 4: Background Isolate Best Match to a Preexisting Genome Assembly**

| Species | Best Match to a Preexisting Genome Assembly | Strain Name in Preexisting Assembly | Percent Similarity* | Plasmids? |
|---|---|---|---|---|
| *Brevibacterium linens* | Assembly_BLIN9172 | *Brevibacterium linens* Strain ATCC 9172 | 99.4% | Unknown |
| *Corynebacterium matruchotii* | ASM17537v1 | *Corynebacterium matruchotii* Strain ATCC 14266 | 99.7% | Unknown |
| *Cutibacterium acnes* | ASM381278v1 | *Cutibacterium acnes* Strain FDAARGOS_503 | 99.3% | No |
| *Escherichia coli* | ASM25814v1 | *Escherichia coli* Strain W | 97.9% | Yes (2) |
| *Lactobacillus gasseri* | ASM1442v1 | *Lactobacillus gasseri* Strain ATCC 33323 | 96.3% | No |
| *Micrococcus luteus* | ASM609441v1 | *Micrococcus luteus* Strain ATCC 4698 | 92.7% | No |
| *Staphylococcus epidermidis* | ASM764v1 | *Staphylococcus epidermidis* Strain ATCC 12228 | 97.4% | Yes (6) |
| *Streptococcus pyogenes* | ASM74301v1 | *Streptococcus pyogenes* Strain ATCC 19615 | 97.8% | No |

*The percent similarity metric is the "genome fraction" reported by the QUAST -R after aligning our SPAdes assembly to the best preexisting reference assembly.

The best preexisting reference assemblies for *Brevibacterium linens*, *Lactobacillus gasseri*, *Micrococcus luteus*, *Staphylococcus epidermidis*, and *Streptococcus pyogenes* have exact strain names matching to those in the CDC AR Isolate Bank. *Cutibacterium acnes* and *Escherichia coli* also have exact strain name matches after reviewing metadata in their respective BioSample (www.ncbi.nlm.nih.gov/biosample/SAMN10163179/) and ATCC documentation (www.atcc.org/products/all/9637.aspx). *Corynebacterium matruchotii* matched to an assembly for ATCC 14266 rather than the expected 14265. An assembly for ATCC 14265 was not available, but its ATCC 14266 near neighbor does have an available preexisting assembly, so that best match makes sense in this case.

The detailed work in characterizing the pathogen and background isolate genomes allowed for the creation of custom taxonomic databases for Bowtie2 read mapping and Mash screen containment calculations, as well as a custom gene database for use with ABRicate, prokka, and Bowtie2. This encompassed pathogen and background organism genes predicted *de novo* by prokka within the SPAdes assemblies and genes annotated within the best preexisting reference assemblies. Supplementary Table 5 shows the number of genes annotated by prokka in our SPAdes isolate assemblies compared to those in the reference assemblies, and all of this information was compiled into a custom gene database for the organisms in our study.

**Supplementary Table 5: Summary of Custom Gene Database Content**

| Organism Type | Species | Direct Match Annotations* | Unique Prokka Annotations | Unique Reference Annotations | Total Number of Genes |
|---|---|---|---|---|---|
| Background | *Brevibacterium linens* | 89.47% | 4.43% | 6.10% | 3,723 |
| Background | *Cutibacterium acnes* | 83.49% | 5.16% | 11.35% | 2,598 |
| Background | *Corynebacterium matruchotii* | 92.28% | 4.37% | 3.35% | 2,629 |
| Background | *Escherichia coli* | 88.15% | 1.53% | 10.32% | 5,155 |
| Background | *Lactobacillus gasseri* | 93.61% | 1.47% | 4.91% | 1,832 |
| Background | *Micrococcus luteus* | 83.08% | 5.29% | 11.63% | 2,477 |
| Background | *Staphylococcus epidermidis* | 87.09% | 2.77% | 10.14% | 2,563 |
| Background | *Streptococcus pyogenes* | 89.74% | 3.23% | 7.04% | 1,890 |
| Background Total | 8 Background Isolates | 88.21% | 3.42% | 8.37% | 22,867 |
| Pathogen | *Clostridioides difficile* | 93.42% | 1.69% | 4.89% | 3,846 |
| Pathogen | *Enterobacter cloacae* | 88.14% | 7.13% | 4.74% | 5,866 |
| Pathogen | *Acinetobacter baumannii* | 92.23% | 2.06% | 5.71% | 4,130 |
| Pathogen | *Enterococcus faecium* | 77.27% | 6.39% | 16.33% | 3,159 |
| Pathogen | *Klebsiella aerogenes* | 90.15% | 1.95% | 7.89% | 5,525 |
| Pathogen | *Klebsiella pneumoniae* | 87.29% | 2.07% | 10.64% | 6,224 |
| Pathogen | *Pseudomonas aeruginosa* | 89.32% | 2.59% | 8.09% | 6,899 |
| Pathogen | *Staphylococcus aureus* | 86.28% | 2.84% | 10.88% | 3,061 |
| Pathogen Total | 8 Pathogen Isolates | 88.43% | 3.29% | 8.28% | 38,710 |

*Direct match annotations are those applied by prokka to our SPAdes assemblies that came directly from the best preexisting reference assembly.

A custom gene database was generated to curate the gene content specific to the 16 background and pathogenic organisms used in this project. To generate the custom gene database, each assembled isolate was annotated through prokka against a reference annotation (see the best preexisting reference assemblies in Supplementary Tables 3 and 4) using default parameters. The resulting prokka annotations for each isolate (i.e., the *.fna files comprised of all the predicted genes), were combined with the remaining annotations of the reference isolate that were not identified by prokka. These retained all possible genes from the isolates in our custom database. All gene predictions retained a unique annotation ID, wherein the source of each prediction was indicated according to the following notation: 1) if prokka identified a prediction within our SPAdes assembly that directly matched an annotation from the best preexisting reference assembly, then annotations were labeled as "*{Isolate_name}_ISOATEDB*"; 2) if prokka identified a prediction within our SPAdes assembly that was not identified in the reference, then annotations were labeled as "*{Isolate_name}_PROKKADB*"; and 3) if prokka did not identify a prediction within our SPAdes assembly that was annotated in the reference, these annotations were labeled as *"{Isolate_name }_REFERENCEDB"*.

The final custom reference database was comprised of 61,577 total predictions across all 16 organisms. A breakdown of the source for each prediction across isolates are listed in Supplementary Table 5. The majority of the gene annotations had direct matches with the reference assembly. *E. faecium* showed the fewest number of direct matches, with only ~77% of the annotations matching to the reference. Annotations that were unique to prokka (i.e. not found in the reference) were generally low (<8%) for all isolates, with the majority of these labeled by prokka as hypothetical proteins. Additionally, <4% of the total predictions in the custom database were identified as pseudogenes. In total, there were 11,311 unique product names in the database when compared with the reference annotations, and 6,157 of these were identified in pathogen isolates, but not in the background isolates. These unique predictions were further investigated and described the function prediction analyses using both mapped reads and assembled contigs. Supplementary Figure 2 shows the shared genome and gene content of the genomes analyzed in this study.

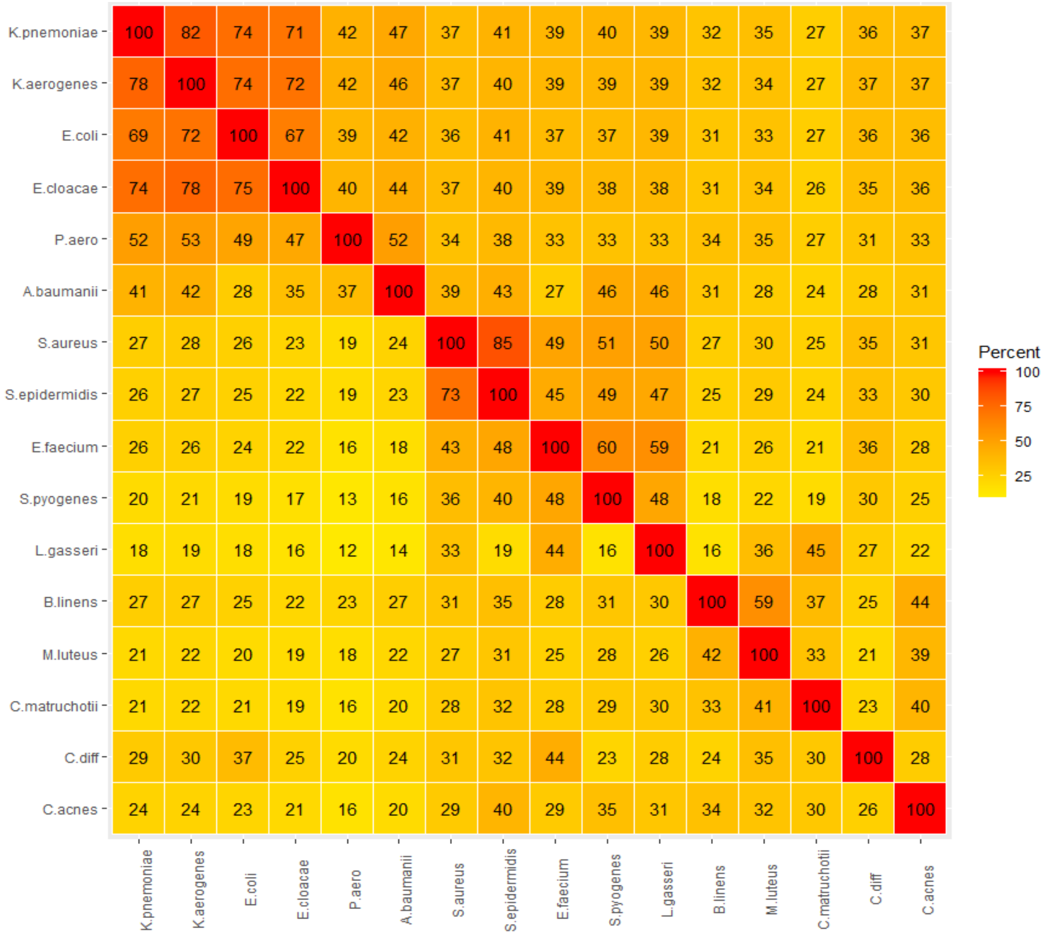

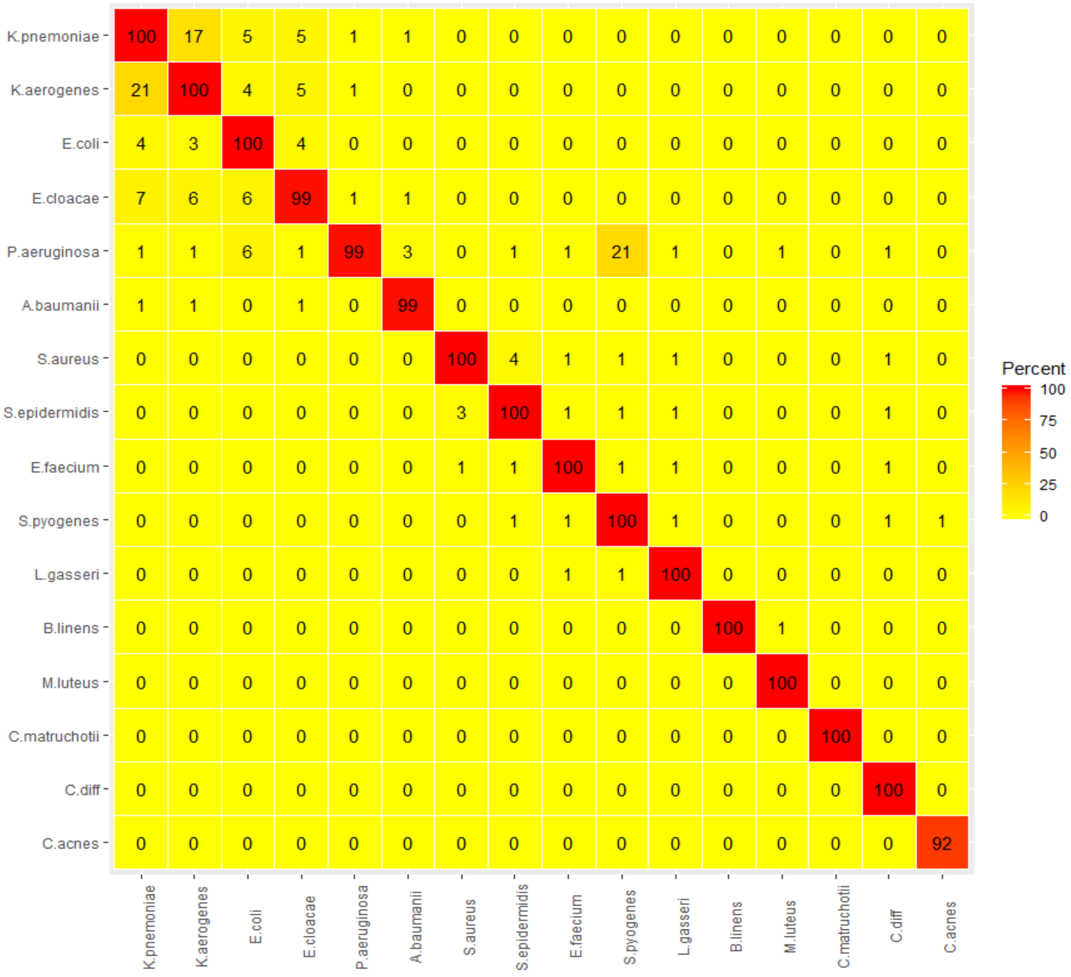


**Supplementary Figure 2. Shared Genic and Total Sequence Content.** A pairwise comparison of all genomes shows shared gene content (top) and shared genome content (bottom). The shared gene content was calculated with prokka reference-guided annotations, and the shared genome content was calculated by pairwise read mapping of all quality filtered reads to each isolate genome. It is clear from this analysis that the overall shared sequence content increases when looking at shared coding regions (left) compared to all coding and non-coding regions of the genomes (right). An increase in shared gene content is particularly evident among *Klebsiella pneumoniae, Klebsiella aerogenes*, *Escherichia coli,* and *Enterobacter cloacae.*

Supplementary Table 6 lists the antibiotic resistance expected for the ESKAPE+C pathogens from https://wwwn.cdc.gov/ARIsolateBank/Search. (Accessed: 19th November 2018)

**Supplementary Table 6: ESKAPE+C Pathogens and Corresponding Antibiotic Resistance**

| Pathogen | Antibiotic Resistance | Molecular Mechanisms of Resistance |
|---|---|---|
| *Acinetobacter baumannii* CDC 275 | Amikacin, Ampicillin/sulbactam, Cefepime, Cefotaxime, Ceftazidime, Ceftriazone, Ciprofloxacin, Doripenem, Gentamicin, Imipenem, Levofloxacin, Meropenem, Piperacillin/tazobactam, Teracycline, Tobramycin | **Aminoglycoside**: aph(3')-Ic, armA, strA, strB; **Beta-lactam**: ADC-25, OXA-23, OXA-66, TEM-1D; **Macrolide-Lincosamide-Streptogramin**: mph(E), msr(E); **Sulfonamide**: sul2 |
| *Clostridium difficile* endospores ATCC 43598 | N/A | N/A |
| *E. aerogenes* CDC 161 | Ampicillin, Ampicillin/sulbactam, Aztreonam, Cefazolin, Cefepime, Cefotaxime, Cefoxitin, Ceftazidime, Ceftazidime/avibactam, Ceftolozane/tazobactam, Ceftriaxone, Doripenem, Ertapenem, Gentamicin, Piperacillin/Tazobactam, Tobramycin | **Aminoglycoside**: aac(3)-IId, aac(6')Ib-cr, strA, strB; **Beta-lactam**: IMP-4, OXA-1, SFO-1, TEM-1B; **Sulfonamide**: sul1; **Phenicol**: catB3; |
| *Enterobacter cloacae* CDC 365 | Ampicillin, Ampicillin/sulbactam, Aztreonam, Cefazolin, Cefepime, Cefotaxime, Cefoxitin, Ceftazidime, Ceftolozane/tazobactam, Ceftriaxone, Ciprofloxacin, Doripenem, Ertapenem, Gentamicin, Imipenem, Levofloxacin, Meropenem, Piperacillin/tazobactam, Tetracycline, Tobramycin, Trimethoprim/sulfamethoxazole | **Beta-lactam**: ACT-type |
| *Enterococcus faecium* CDC 579 | Ampicillin, Doxycycline, High-Level Gentamicin, High-Level Streptomycin, Penicillin, Quinupristin/dalfopristin, Rifampin | **Aminoglycoside**: ant(6)-Ia, aph(2'')-Id, aph(3')-III; **Macrolide-Lincosamide-Streptogramin**: lnu(B); **Tetracycline**: tet(U) |
| *Klebsiella pneumoniae* CDC 139 | Amikacin, Ampicillin, Ampicillin/sulbactam, Aztreonam, Cefazolin, Cefepime, Cefotaxime, Cefoxitin, Ceftazidime, Ceftazidime/avibactam, Ceftolozane/tazobactam, Ceftriaxone, Ciprofloxacin, Doripenem, Ertapenem, Gentamicin, Impipenem, Levofloxacin, Meropenem, Piperacillin/tazobactam, Tetracycline, Tobramycin, Trimethoprim/sulfamethoxazole | **Aminoglycoside**: aac(6')-IIa, armA, strA, strB; **Beta-lactam**: NDM-1, CMY-4, CTX-M-15, OXA-10, SHV-11; **Macrolide-Lincosamide-Streptogramin**: mph(E), msr(E); **Fosfomycin**: fosA; **Sulfonamide**: sul1, sul2; **Phenicol**: cmlA1; **Rifampicin**: ARR-3; **Trimethoprim**: dfrA1 |
| *Pseudomonas aeruginosa* CDC 230 | Amikacin, Cefepime, Ceftazidime, Ciprofloxacin, Doripenem, Gentamicin, Imipenem, Levofloxacin, Meropenem, Piperacillin/tazobactam, Tobramycin | **Aminoglycoside**: aac(3)-Id, aadA2; **Beta-lactam**: VIM-2, OXA-4, OXA-50, PAO; **Phenicol**: cmlA1; **Tetracycline**: tet(G); **Trimethoprim**: dfrB5 |
| *Staphylococcus aureus* CDC 219 | Cefoxitin, Clindamycin, Erythromycin, Gentamicin, Levofloxacin, Oxacillin, Penicillin, Rifampin, Tetracycline | **Aminoglycoside**: aac(6')-aph(2''), aadD, spc; **Beta-lactam**: mecA; **Tetracycline**: tet(M); **Macrolide-Lincosamide-Streptogramin**: erm(A) |

**Supplementary Table 7: Bioinformatics Tools**

| Software | Description | Command | Publication |
|---|---|---|---|
| Trimmomatic | Removal of sequence adapters and quality control read trimming with Trimmomatic http://www.usadellab.org/cms/?page=trimmomatic | `trimmomatic PE {input_1.fq.gz} {input_2.fq.gz} {output_pe_1.fq.gz} {output_se_1} {output_pe_2.fq.gz} {output_se_2} ILLUMINACLIP:illumina-adapters.fa:2:40:15 LEADING:{trim_value} TRAILING:{trim_value} SLIDINGWINDOW:4:{trim_value} MINLEN:25 -trimlog {output_log}` | Bolger, Lohse, and Usadel 2014 |
| FastQC | Production of quality control reports for sequence reads with FastQC before and after trimming https://www.bioinformatics.babraham.ac.uk/projects/fastqc/ | `fastqc {input_filename.fq.gz} -o {output_directory_name_fastqc}` | Andrews 2010 |
| MultiQC | Aggregation and visualization of FastQC read quality statistics https://multiqc.info/ | `multiqc {inputs_fastqc.zip} -n {output_multiqc_fastqc_report_name }` | Ewels et al. 2016 |
| SPades | Assembly of bacterial isolates with SPAdes http://cab.spbu.ru/software/spades/ | `spades.py -k 21,33,55,77 --careful -1 {input_1.fq.gz} -2 {input_2.fq.gz} -o {output_name}` | Bankevich et al. 2012 |
| metaSPAdes | Metagenome assembly with metaSPAdes http://cab.spbu.ru/software/spades/ | `metaspades.py -1 {input_1.fq.gz} -2 {input_1.fq.gz} -o {output_name}` | Bankevich et al. 2012 |
| rnaSPAdes | Transcriptome assembly with rnaSPAdes http://cab.spbu.ru/software/spades/ | `rnaspades.py -1 {input_1.fq.gz} -2 {input_1.fq.gz} -o {output_directory_name}` | Bankevich et al. 2012 |
| QUAST | Assembly evaluation with QUAST http://bioinf.spbau.ru/quast | `quast.py {input_contigs.fasta} -R {reference_genome} -o {output_quast_directory}` | Gurevich et al. 2013 |
| MultiQC | Aggregation and visualization of QUAST assembly statistics https://multiqc.info/ | `multiqc {inputs_quast_report.tsv} -n {output_multiqc_quast_report_name} -o {output_name}` | Ewels et al. 2016 |

| Software | Description | Command | Publication |
|---|---|---|---|
| Bowtie2 | Alignment of sequences to reference http://bowtie-bio.sourceforge.net/bowtie2/manual.shtml | Indexed the gene sequences using Bowtie2 with this command: `find *fasta *fna \| parallel bowtie2-build {} {}` Ran this command for a set of trimmed sequences in the same directory as indexed sequences: `find *fasta \| parallel "bowtie2 --threads 4 -x {} -1 {sample}_trim30_1.fastq.gz -2 {sample}_trim30_2.fastq.gz` | Langmead and Salzberg, 2012 |
| SAMtools | Viewing and sorting of the aligned reads http://www.htslib.org/doc/samtools.html | `samtools view -bS \| samtools sort > {/.}.{sample}.sorted.bam" && find *.sorted.bam` | Li et al., 2009 |
| Qualimap2 | Visualize and calculate alignment statistics http://qualimap.conesalab.org/doc_html/index.html | `parallel -j4 "qualimap bamqc --java-mem-size=20G -nt 12 -bam {} -outdir {sample}-{/.}" && mkdir qualimap-{sample} && mv {sample}* qualimap-{sample}/ && multiqc --title {sample}-qualimap-bowtie2 --force qualimap-{sample}` | Okonechnikov et al., 2016 |
| Mash | Distance estimation between an isolate and reference genome https://mash.readthedocs.io/en/latest/tutorials.html | `mash dist {reference_db.msh} {input_reads.fastq} > {output.distances.tab}` | Ondov et al. 2016 |
| Mash Screen | Screening of a metagenome or metatranscriptome for the containment of reference genomes https://mash.readthedocs.io/en/latest/tutorials.html | `mash screen {reference_db.msh} {input_reads.fastq} > {output.screen.tab}` | Ondov et al. 2019 |
| Prokka | Prokaryotic *de novo* gene predictions from isolate contigs https://github.com/tseemann/prokka | `prokka {input_contigs.fa} --rnammer --outdir {output_directory_name} --prefix {output_filename_prefix}` | Seemann 2014 |

| Software | Description | Command | Publication |
|---|---|---|---|
| Prokka | Prokaryotic *de novo* gene predictions from metagenome contigs https://github.com/tseemann/prokka | `prokka {input_contigs.fa} --metagenome --evalue 1e-06 --notrna --rnammer --outdir {output_directory_name} --prefix {output_filename_prefix}` | Seemann 2014 |
| ABRicate | Gene identifications from contigs https://github.com/tseemann/abricate | `abricate {input_contigs.fa} --csv --db {database_name} > {output_name.csv}` | Zankari et al. 2012 |
| Mauve | Multiple Genome Aligner http://darlinglab.org/mauve/user-guide/progressivemauve.html | `Java -Xmx500m -cp {path/to/Mauve.jar} org.gel.mauve.contigs.ContigOrderer -output {output_directory_name} -ref {reference_genome.gbk} -draft {draft_contigs.fasta}`<br><br>`progressiveMauve –output={output_filename} {each_genome_to_align.gbk} {rearranged_draft_contigs.fasta}` | Darling et al. 2010 |

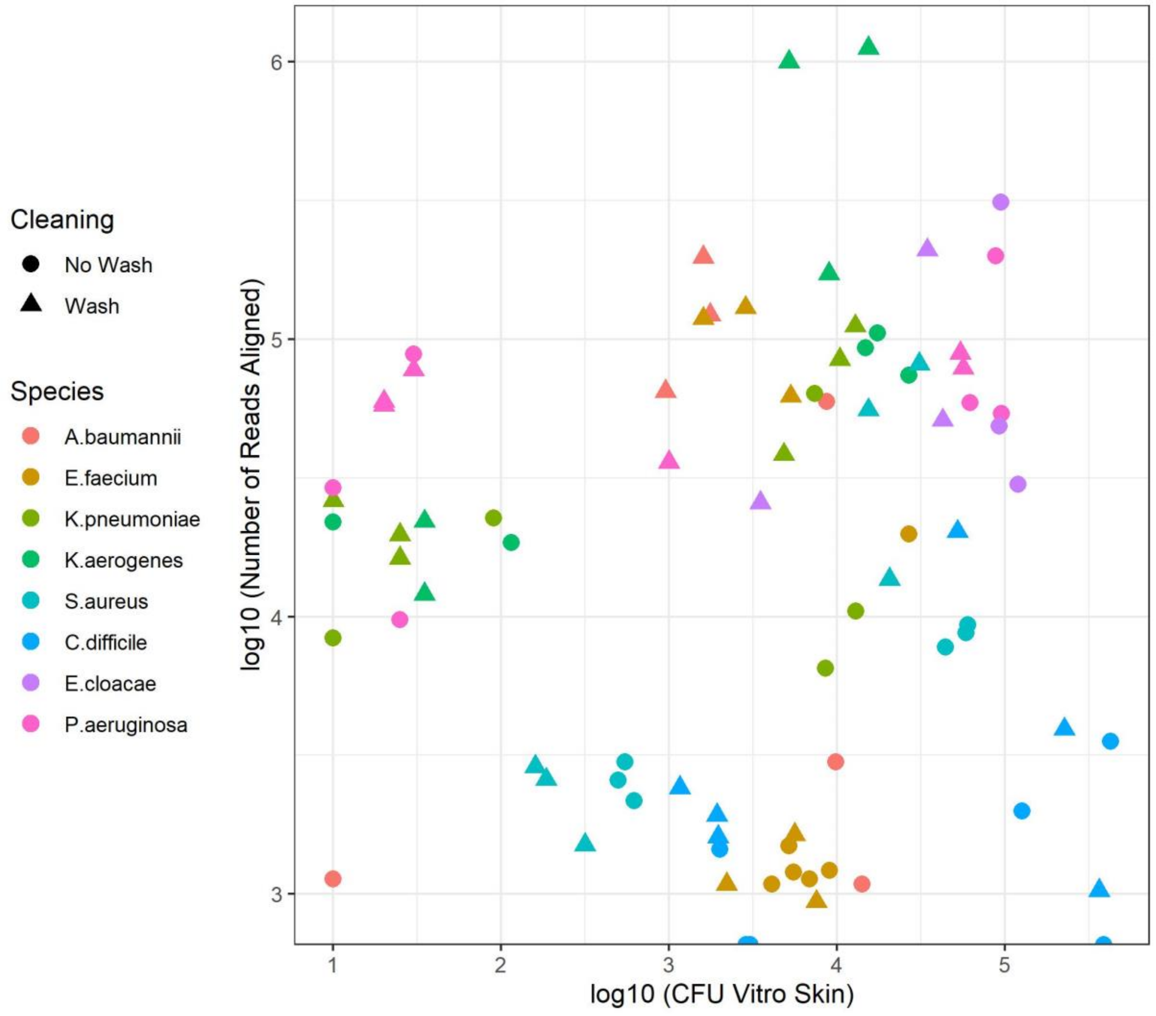


**Supplementary Figure 3. CFU vs. Mapped Reads in Direct Transfer Scenarios.** There was not an observable correlation between mapped reads and CFU counts.

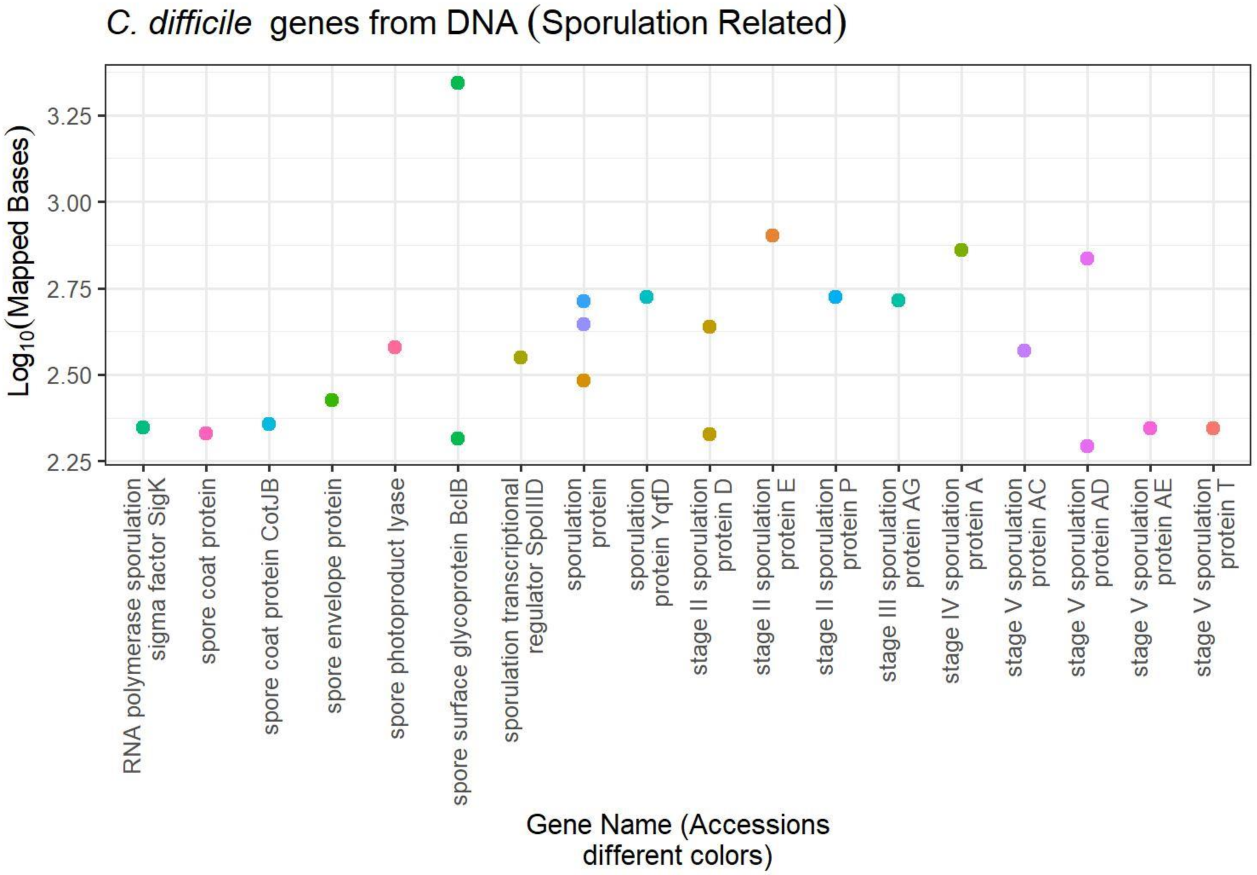


**Supplementary Figure 4. Sporulation-Related Genes Identified in Metagenomes from *C. difficile*.** Sporulation related genes from *C. difficile* were detected by read mapping in metagenomics direct transfer scenarios, despite being below detection by culture techniques. The spore-related genes represent a subset of the 621 accessions representing 337 unique protein names that were detected. The sporulation subset shown were only identified within the direct transfers scenarios during the handwashing simulation.

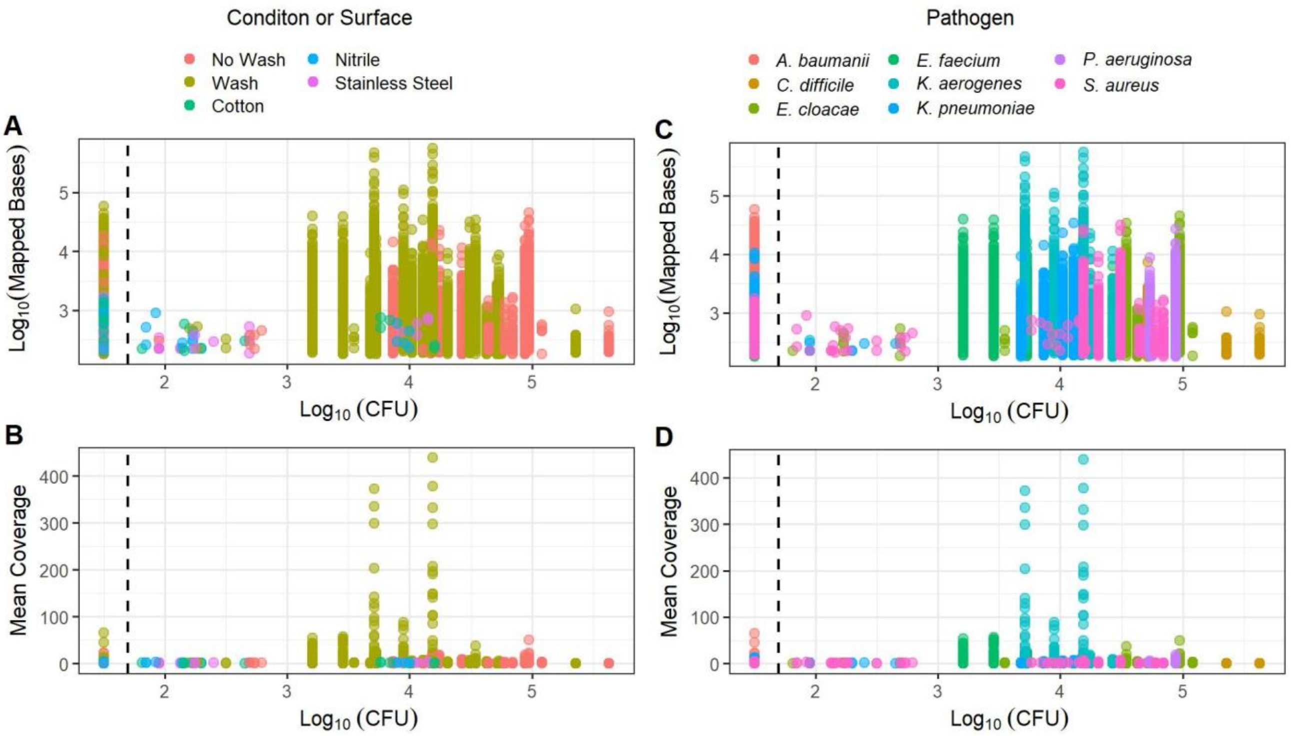


**Supplementary Figure 5. Indirect Scenario Pathogen CFU vs Metagenome Reads Mapped to Genes.** Comparison of mapped sequences to genomes ($Log_{10}$ Mapped Bases or Mean Coverage) and culturing of pathogens ($Log_{10}$ CFU) from contact scenarios. CFU values below the detection limit (dotted line, 50 CFU, ~ $Log_{10}$ 1.7) were converted to $Log_{10}$ 1.5 to appear on figure.

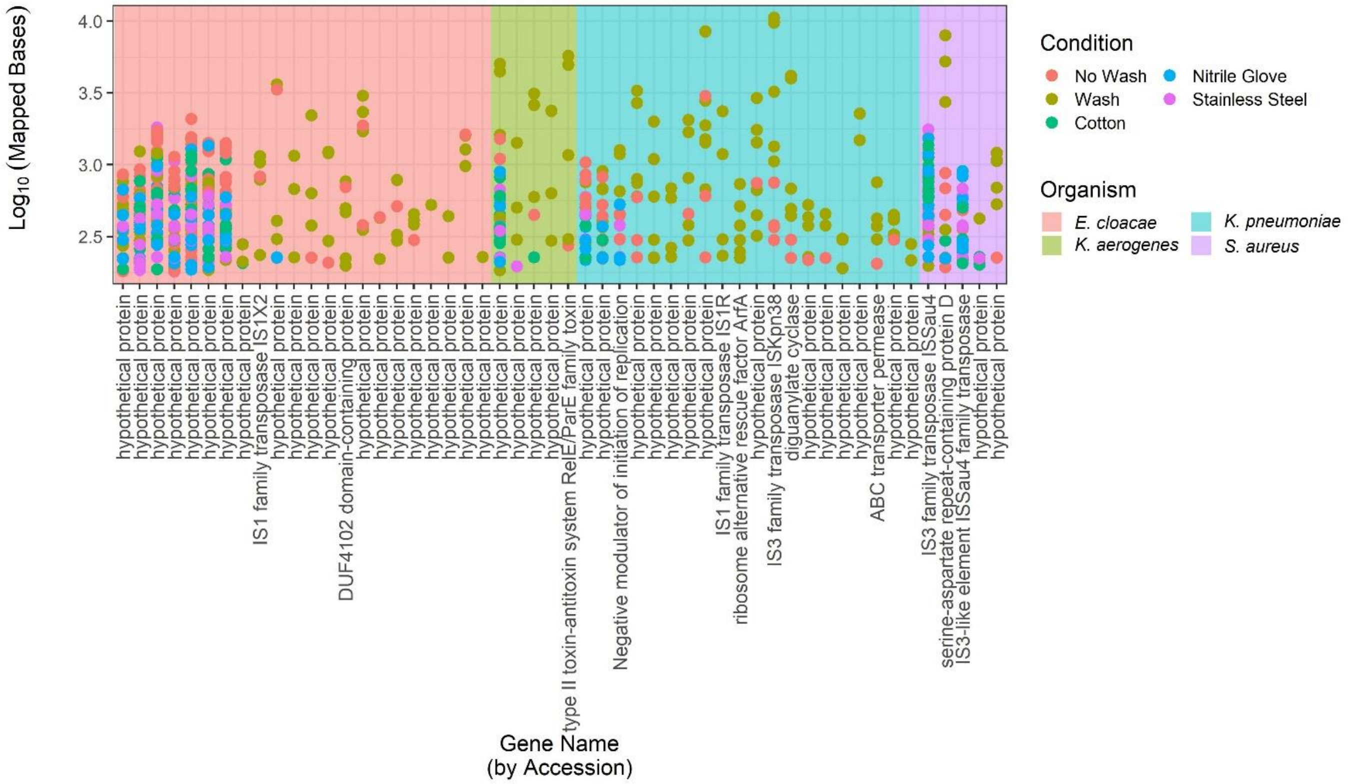


**Supplementary Figure 6. DNA Reads Mapped to Resilient False Positive Genes.** After the remove of shared genes and the use of a 180 bp mapped bases cutoff, 52 accessions remained that correlated with pathogens used outside of the mixtures that were sequenced. These 52 accessions were removed from further analysis.

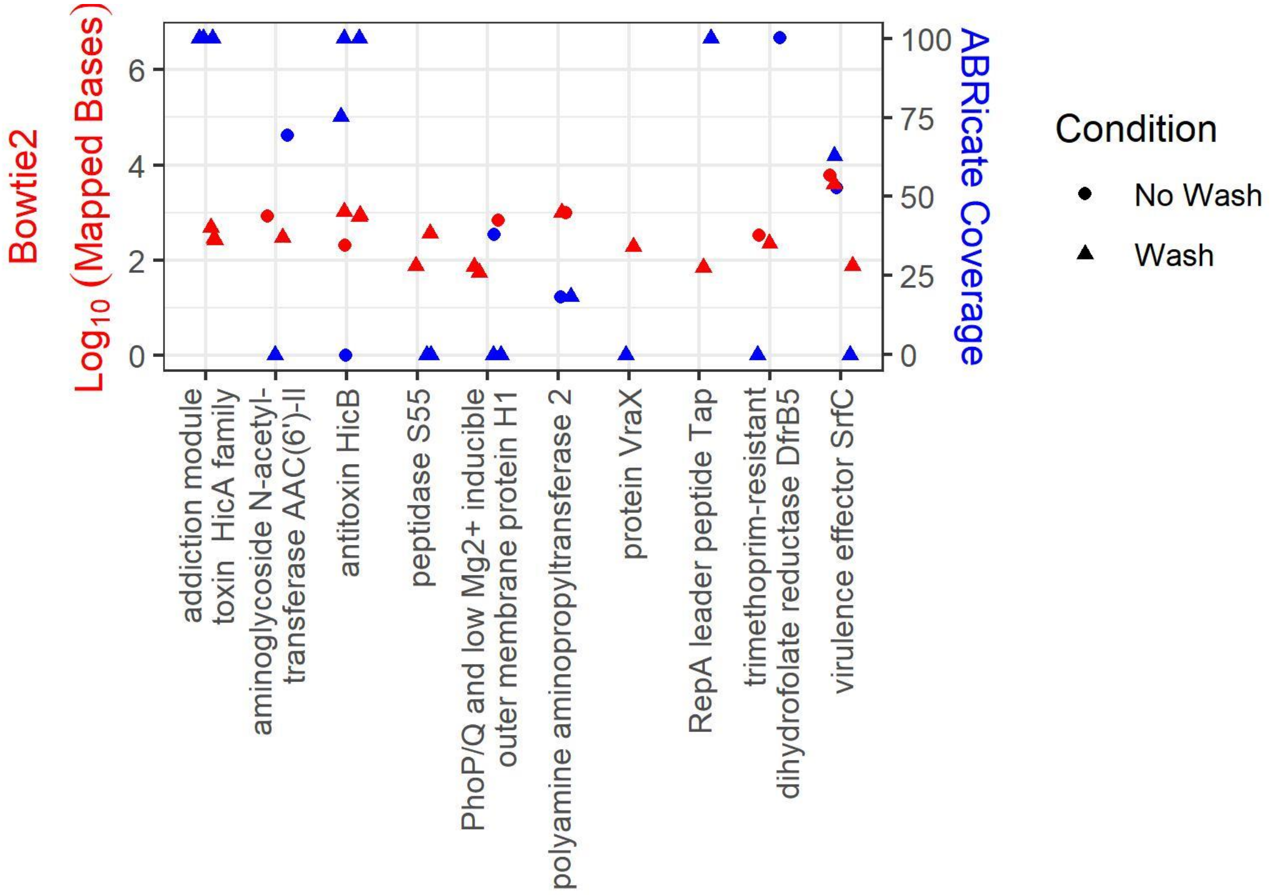


**Supplementary Figure 7. ABRicate vs. Bowtie2 Gene Detection.** Comparison of mapping of reads using Bowtie2 and annotation of assembled reads using ABRicate from contact scenarios with shared and known false positive accessions within Bowtie2 read mapping. Remaining accessions after elimination included direct contact scenarios of high inoculum what were simulated wash and no wash.

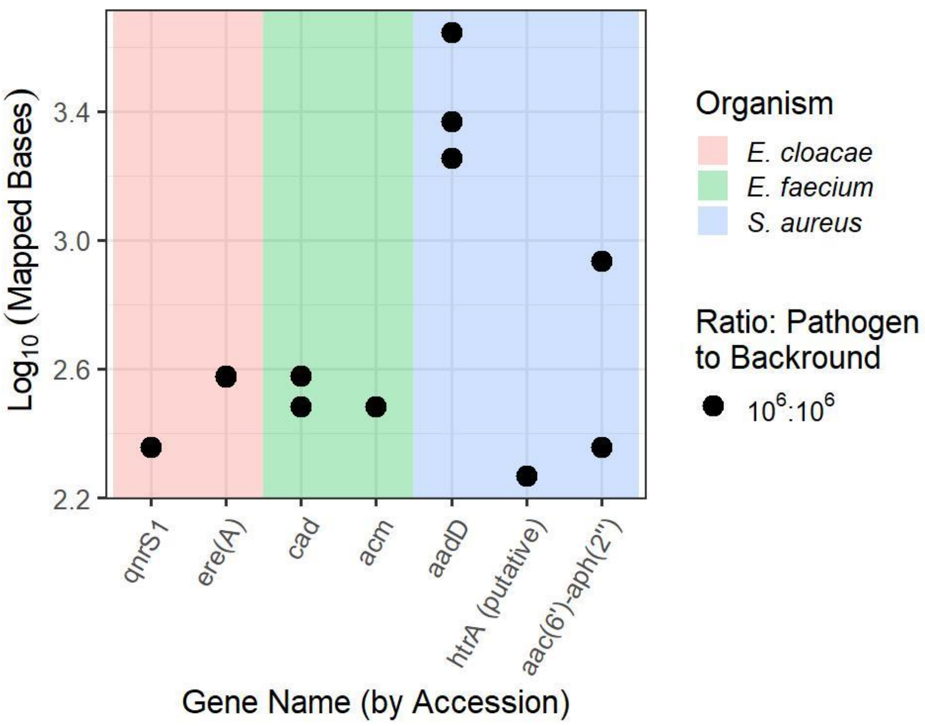


**Supplementary Figure 8. RNA Reads Mapped to AMR Genes.** Metatranscriptomic sequences derived from RNA in spike-ins were mapped to genes related to antimicrobial resistance (AMR) and virulence. The two genes detected from *E. cloacae* within the metagenome contact scenarios were also detected with RNA transcripts suggesting active expression, while the majority of genes were not detected at the transcriptional level which may suggest lack of expression or transcripts were below the detection limit.

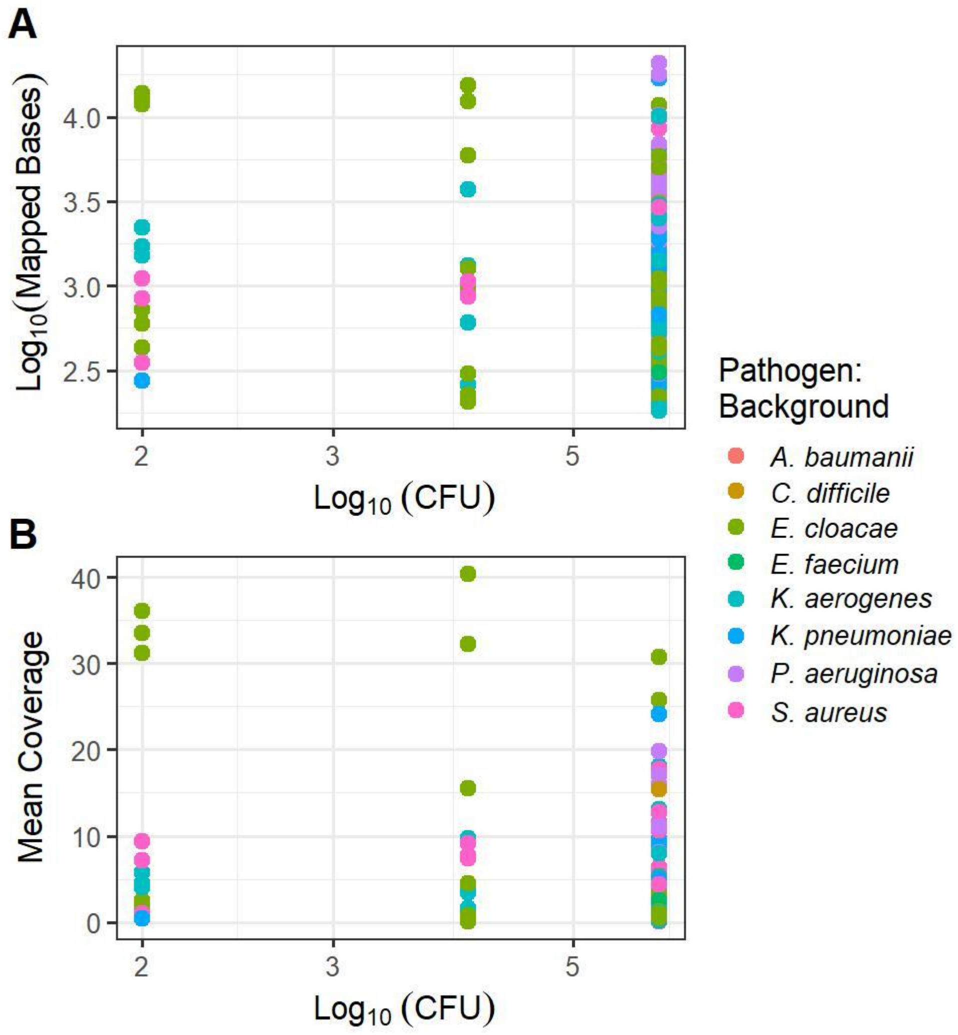


**Supplementary Figure 9. RNA Reads Mapped to Genes vs CFU.** Comparison of mapped sequences, from RNA extracted from spike-ins of pathogens to background organisms, to the collective genomes utilizing bowtie2 ($Log_{10}$ Mapped Bases or Mean Coverage) and estimated CFU ($Log_{10}$ CFU). Mapped results were filtered as contact scenarios to remove shared genes, mapped base pair cutoff of 180, and additional known false positive accessions. The relationship of mapped bases and mean coverage in relation to estimated CFU from the spike-ins pf pathogens to background organisms were dependent on the organism. Different genes specific to an organism within the mixture demonstrated variability with some within the organisms showing a positive direct linear relationship with CFU, while others appeared static with increasing estimated CFU.
